# Mechanistic modeling and machine learning identifies optimum radiotherapy schedules to prevent treatment-induced metastasis

**DOI:** 10.1101/2025.07.02.662373

**Authors:** Christopher Graser, Zhi Zhou, Manuel Schürch, Graydon Moorhead, Justin Dean, Cindy Lin, Jamie Dean, David Kozono, Galit Lahav, Franziska Michor

**Author notes:** These authors contributed equally to this work (Christopher Graser, Zhi Zhou).

## Abstract

Lung cancer patients often experience increased metastasis formation after radiotherapy. However, it is incompletely understood whether radiation affects the migratory behavior of tumor cells and how altered radiotherapy schedules might mitigate this risk. To address these questions, we performed live-cell microscopy experiments to profile changes in cell migration during radiation across 12 cancer cell lines and developed a predictive computational modeling platform describing tumor volume and dissemination during radiotherapy. Using this platform, we identified optimal fractionation schedules and then performed extensive *in silico* clinical trials, establishing that our optimized schedules substantially reduce metastatic seeding relative to the standard of care schedule. Training transformer models on the *in silico* clinical trial data enabled us to recover mechanistic parameters with high accuracy, demonstrating that the features determining optimal radiotherapy can be inferred from longitudinal tumor data. Our integrative predictive approach enables the rational design of optimum clinical trials across indications.

## Introduction

Non–small cell lung cancer (NSCLC) accounts for approximately 80% of lung cancer cases and remains the leading cause of cancer-related mortality worldwide (Siegel et al., 2024). Radiotherapy plays a crucial role in the treatment of NSCLC, especially for patients with inoperable tumors, incompletely resected tumors, or recurrent disease following surgery (Durante and Loeffler, 2010; Baskar et al., 2014). However, recurrences after radiotherapy are often associated with distant metastasis. Accumulating evidence suggests that radiotherapy may inadvertently promote metastatic spread by promoting migratory and invasive features in NSCLC cells (Jung et al., 2007; Ho et al., 2010; Zhou et al., 2011). As early as 1949, Kaplan and Murphy (1949) observed increased lung metastases in tumor-bearing mice when treated with ionizing radiation compared to sham-treated mice. Martin et al. (2014) reported that radiotherapy increases the number of circulating tumor cells (CTCs) in NSCLC patients during the course of treatment, and Frick et al. (2020) showed that persistently high levels of CTCs during stereotactic radiotherapy of NSCLC are associated with treatment failure due to distant metastases. Several mechanisms related to different steps of the metastatic seeding process have been proposed to explain this phenomenon, including radiation-induced changes in cell-cell adhesion, cell motility, interactions with the stromal environment, and the propensity of cells to undergo an epithelial to mesenchymal transition (Moncharmont et al., 2014). Among these factors, cell motility appears to be a key component of the metastatic cascade (Stuelten et al., 2018).

The phenomenon of potentially radiation-induced metastasis is not unique to lung cancer (Moncharmont et al., 2014; Blyth et al., 2018), but there are some differences in the mechanisms that have been implicated in local tissue invasion versus distant spread across cancer types. For instance, Paquette et al. (2011) demonstrated that irradiating fibroblasts prior to co-culturing them with unirradiated breast cancer cells significantly increases tumor cell invasiveness via increased fibroblast cyclooxygenase-2 expression; however movement in the absence of extracellular matrix proteins was not affected. Conversely, Bouchard et al. (2017) showed in a mouse model of triple negative breast cancer that irradiation both increased local invasiveness and the frequency of lung metastases and that both effects were driven by induction of interleukin-1β expression. Notably, studies in different cancer types, or even different cell lines from similar cancer types, found heterogeneous effects of irradiation on metastatic activity (Charmont et al., 2014; Blyth et al., 2018). For instance, Woods et al. (2015) demonstrated that radiotherapy was associated with an increased risk of metastatic seeding for alveolar rhabdomyosarcoma xenografts in mice, whereas for embryonal rhabdomyosarcoma xenografts, exposure to radiotherapy was found to be associated with a decreased risk of metastatic seeding. Thus the connection between radiotherapy and cell migration and metastasis is heterogeneous across cell types and culture conditions, yet might have important implications for the treatment response of patients receiving radiation therapy.

In the clinic, contemporary standard of care fractionation schedules do not directly take into account potential effects of radiation on metastatic activity. Here, we set out to quantitatively characterize the effect of irradiation on cell migration as a component of the metastatic cascade and to investigate how fractionation schedules can be adapted to mitigate this altered risk. To this end, we performed live-cell imaging experiments investigating changes in cell migratory behavior upon radiation across 12 cell lines from different cancer types (Fig. 1A). We then developed a predictive computational modeling framework to quantitatively describe cell migration in response to radiotherapy based on our experimental findings. This framework allows us to characterize optimal treatment schedules as a function of changes in tumor cell motility in response to radiation (Fig. 1B). Moreover, to quantify the extent to which these schedules may improve treatment outcomes, we performed an *in silico* clinical trial, in which treatment dynamics were simulated for an exemplary patient cohort. The results of this *in silico* clinical trial suggest that the optimized treatment schedules significantly reduce the risk of metastatic seeding associated with increased cell migration compared to standard of care fractionation schedules. Finally, we trained machine learning models on the *in silico* clinical trial data. This approach demonstrated that we can recover mechanistic parameters of the treatment response model with high accuracy, demonstrating that the features which determine optimal radiotherapy schedules can be inferred from longitudinal tumor volume data over short time horizons. This work forms the basis for rational design of personalized clinical strategies aimed at mitigating the risk of radiation-induced metastasis.

**Fig. 1.**
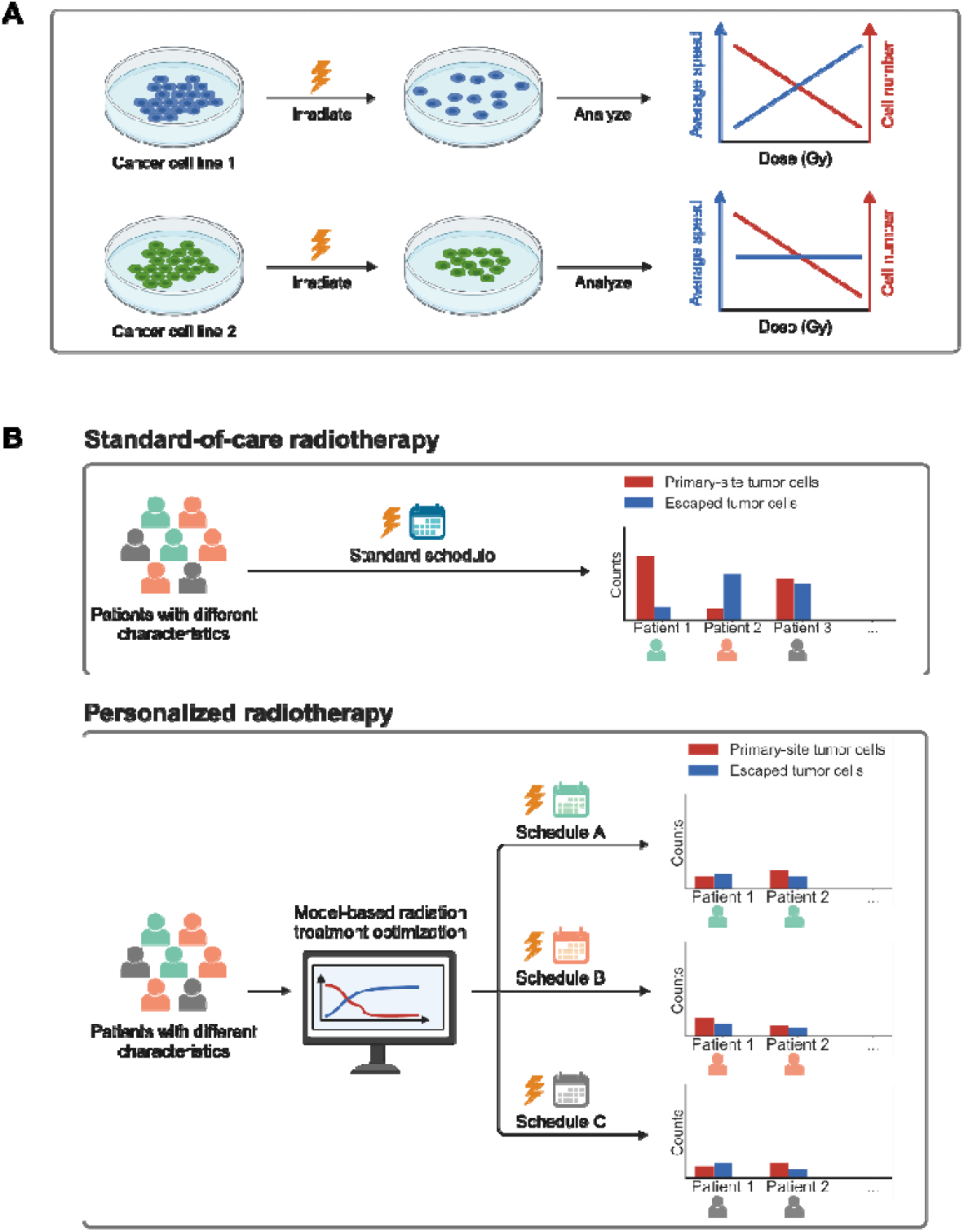
Quantification of the effects of radiation on cell migration enables design of personalized radiotherapy schedules. **A)** Tumor cells may exhibit altered motility in response to radiation, a change that can be inferred from patterns of cell dispersal over time. To quantify the radiation-induced motility changes, we experimentally determined the dispersal patterns of irradiated cells across different tumor cell lines and evaluated the relationship between radiation dose and cell migration speed (indicated by the blue lines), as illustrated in the schematic. **B)** Due to inter-patient heterogeneity in the propensity of tumor cells to escape upon irradiation (with colors representing patients with distinct tumor characteristics), a one-size-fits-all standard schedule may lead to suboptimal outcomes in reducing primary tumor cells (shown in red) and escaped tumor cells (shown in blue). Personalized schedules, derived from patient-specific characteristics through model-based optimization (colors represent patients and their personalized schedules), can improve treatment outcomes in both tumor shrinkage and metastasis prevention (shown in red and blue, respectively).

## Results

### Live-cell experiments reveal heterogeneous dose-speed relationships across cell lines

We analyzed the motion of cells across 12 different cancer cell lines treated with varying radiation doses (1-8Gy) and imaged every 15 minutes for 24 hours using live cell fluorescence microscopy (Fig 2A; Stewart-Ornstein and Lahav, 2017). We estimated the speed of individual cells from the extracted positions of cell centroids, performed linear regression of speed against dose, and calculated the Pearson correlation coefficients (Methods). Across cell lines, we observed substantial variability in the effect of radiation on cell motility, with correlation coefficients varying from-0.44 to 0.47 (Fig. 2B). Among the 12 cell lines, two cell lines (A549 and UO31) exhibited a significant positive correlation between speed and dose with respective Pearson correlation coefficients of *r =* 0.47 and 0.23, suggesting a dose-dependent enhancement in cell motility; six cell lines (NCI-H460, HCT116, MALME3E, MCF7, UACC257, UACC62) showed a significant negative correlation with correlation coefficients ranging from-0.44 to-0.21; the remaining four cell lines showed no correlation. Cell lines pertaining to the same cancer type did not show any particular similarities (see contrasting results for NSCLC cell lines A549 and NCI-H460; and kidney cancer cell lines UO31 and A498). Further analyses (SI1) revealed that the effects of radiation on migratory behavior did not vary with time (SF1,2), were not driven by outliers (SF3), and showed no directional bias (SF4). Interestingly, we found that changes in p53 levels correlate with migratory changes in A549 cells (SI1; SF5,6), implicating p53 as a potential mediator of migratory response (SI1; ST1).

**Fig. 2.**
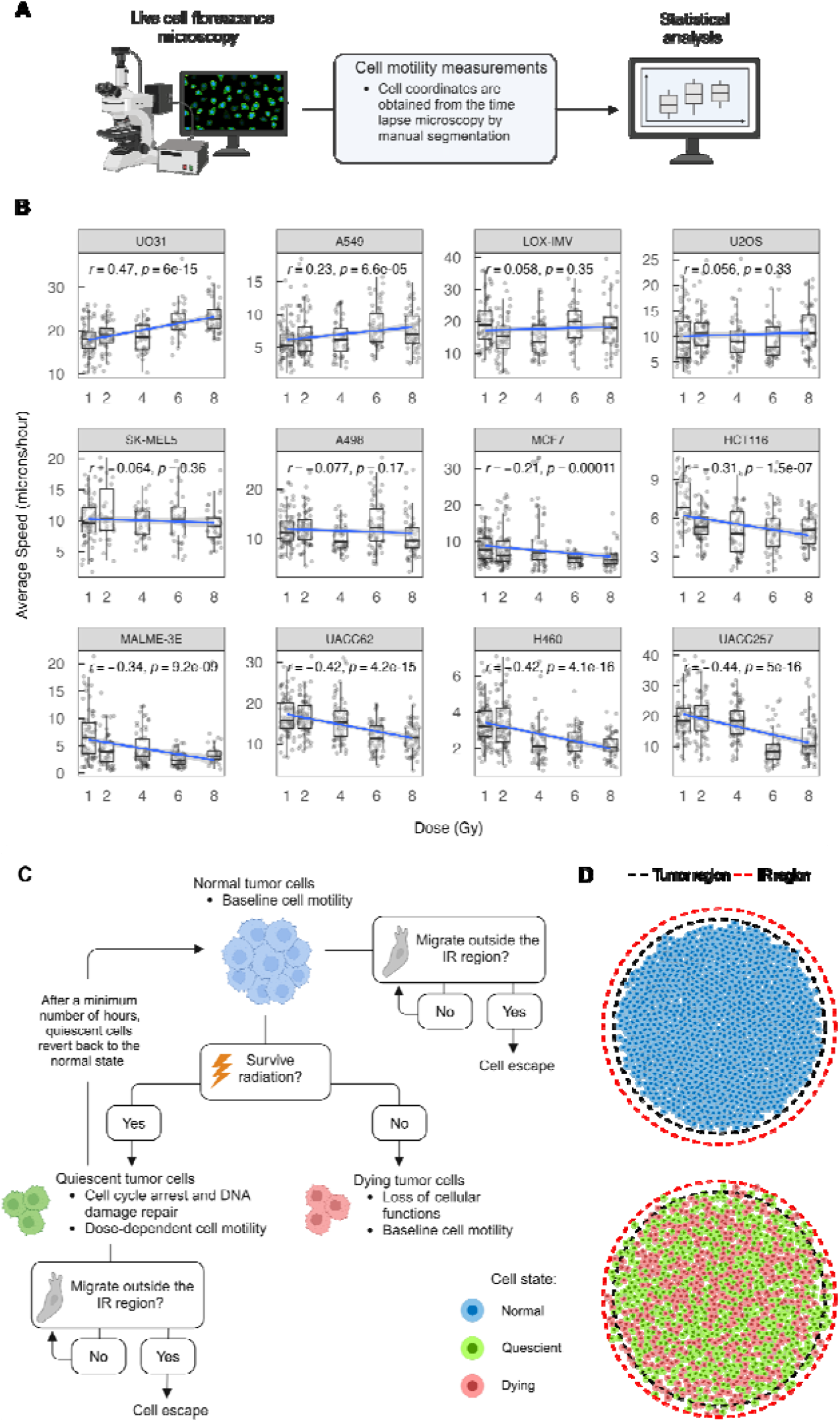
A computational modeling framework for personalized radiation treatment optimization under the risk of treatment-induced metastases. **(A)** Summary of the experimental workflow (see Methods). **(B)** Averaged speeds of cells from 12 cancer cell lines treated with varying radiation doses. Substantial variability is observed in the effect of radiation on cell motility. Correlation between dose and speed was formally tested within each cell line. The Pearson correlation coefficients r and their associated p-values p, as well as the linear regression lines of speed on dose, are displayed. **(C)** Schematic of cellular behaviors in the agent-based simulation of tumor response to radiotherapy. **(D)** Depictions of the spatial locations of tumor cells before and after radiation (IR) treatment in an agent-based simulation. The dashed black circle indicates the boundary of the initial tumor region and the red circle indicates the boundary of the IR region. Tumor cells in the primary tumor region migrate at a baseline speed; after IR, tumor cells become quiescent or die, where quiescent and dying cells exhibit dose-dependent or baseline motility. The risk of metastatic seeding is quantified by the proportion of viable tumor cells that escape the IR region by the end of a radiation schedule.

### A computational model of cell proliferation and movement during radiation

Understanding how the migratory behavior of tumor cells changes in response to radiation may help inform the design of optimal fractionation schedules for radiation therapy, as different dosing regimens may lead to varying risk of metastatic seeding. To identify schedules that optimally reduce both tumor burden and metastatic seeding, we constructed a spatially explicit stochastic agent-based model of tumor cells in response to radiotherapy, where the dose-dependent migration of irradiated tumor cells was parameterized by the cell motility measurements of A549 cells post-radiation (Fig. 2C, Methods). Using this model and considering that the number of tumor cells that escape from the primary tumor site may be positively associated with metastatic risk, we simulated the behaviors and interactions of a collection of individual tumor cells during different radiation fractionation schedules. We then quantified the impact of radiation on the migration of tumor cells and their propensities to seed metastases, characterized by their escape behaviors (Fig. 2A-C).

Tumor response to radiotherapy was modeled using the linear quadratic (LQ) model, which states that the surviving fraction of tumor cells after being irradiated with dose *d* Gy is *S*(*d*) = exp(−*α* × *d* − *β* × *d*^2^ ), where *α* and *β* are tissue-specific radiosensitivity parameters and d denotes the delivered radiation dose (Hall and Giaccia, 2012, Fowler, 2010, and McMahon, 2018). The computational model incorporates cellular movement driven by both random Brownian motion and by cell-cell interactions in a radiation dose-dependent manner. Following irradiation, surviving cells migrate at a speed *u* = *a* + *b* × *d*, where *b* is the dose-speed coefficient and denotes the dependence of a cell’s motility on the radiation dose d (Methods). Cells were counted as escaped if they moved beyond the primary tumor region, defined as a circle with a radius 110% of the tumor’s initial radius (Fig. 2D). This added margin represents the irradiation (IR) margin commonly used in radiotherapy to compensate for tumor localization uncertainties and undetectable microscopic tumor extensions (Burnet et al. 2004, Tamaki et al. 2022). Cells in the primary tumor region remain exposed to radiation, while escaped cells are assumed to sustain little to no radiation damage and are removed from the simulation. The number of escaped tumor cells is associated with the number of invading cells and CTCs, and consequently, the risk of metastatic seeding (Frick et al. 2020).

As stochastic simulations of the full agent-based model are computationally expensive, we also developed a simplified ordinary differential equations (ODE) version of our model (Methods; Fig. 3A). This deterministic model describes the size of a continuous tumor mass, rather than tracking individual cells, and discretizes space into compartments of varying proximity to the tumor IR margin to track movement between but not within these compartments. This ODE version of the model was used to identify optimized schedules, as it allows iteration through a large set of candidate fractionation schedules in a computationally efficient manner. We then validated these optimal schedules with simulations of the agent-based computational model to ensure that the identified schedules indeed significantly outperform standard treatment schedules.

**Fig. 3.**
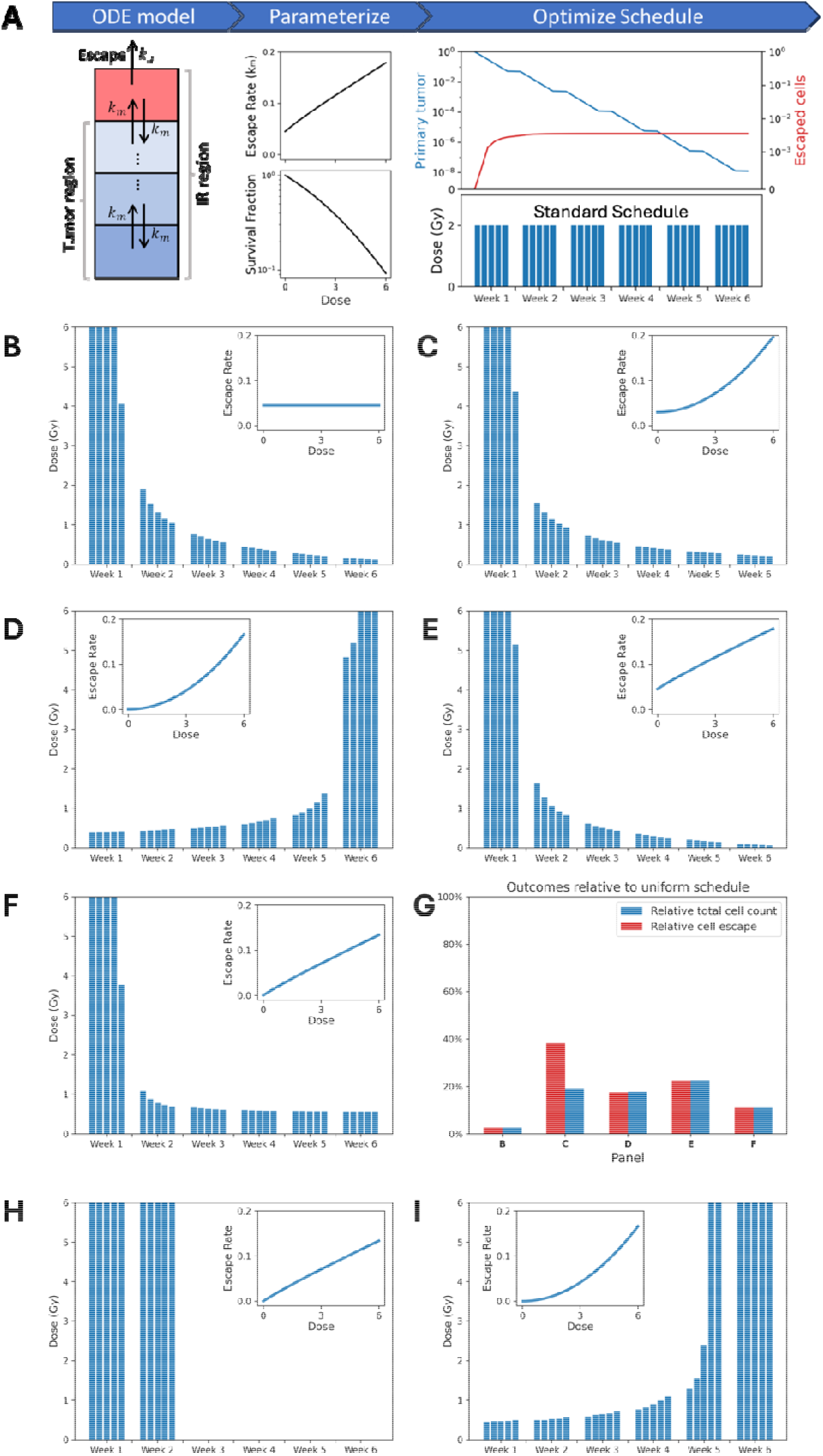
The computational modeling framework identifies optimal fractionation schedules. **(A)** For optimization purposes, we designed an ODE system to approximate the dynamics of the agent-based model. The ODE system was parameterized with a function relating cell survival to the irradiation dose and a function relating cell escape to the irradiation dose. For a given schedule, the system of ODEs identifies a trajectory of the proportion of cells that survive in the primary tumor and the proportion of cells that escape over time. We optimized therapy by choosing schedules that minimize a convex combination of the fraction of escaped cells and the fraction of surviving cells in the primary tumor at the end of treatment (see Methods). **(B)-(F):** Optimal six-week fractionation schedules as a function of the dose-escape relationship (see Methods for parameterization), under the constraints that the total BED is equal to that of the standard of care schedule, and that the maximal dose per day is 6 Gy. **(G)**: Performance of optimized schedules relative to the standard uniform schedule from panel (A). The blue bars indicate the ratio of the total number of cells (escaped and in the primary tumor) at the end of treatment under the optimal fractionation schedule vs. under the uniform schedule, for the dose-escape relationships and respective optimal schedules in panels B through F. The red bars indicate the ratio of the number of cells that have escaped by the end of treatment under the optimal fractionation schedule vs. under the uniform schedule, for the dose-escape relationships and respective optimal schedules in panels (C)-(G). **(H)-(I):** Optimal six-week fractionation schedules for dose-escape relationships as in panels (F) and (D), under the constraint that total absolute doses add up to 60Gy. Under this constraint, the schedule in panel (H) is also optimal for the dose-escape relationships in panels (B),(C) and (E).

### Optimal schedules for different dose-risk relationships

For optimization, schedules were scored based on an equally-weighted sum of the number of viable cells (cycling and quiescent tumor cells) in the primary tumor and the number of viable cells that have escaped by the end of treatment (Methods). The standard radiotherapy dose for stage III NSCLC patients is 60 Gy at 2 Gy per once-daily fraction, which has remained unchanged over the past three decades (Perez et al., 1980; Bradley et al., 2015). Guided by this standard radiation dose, we formulated constraints for our optimization problem. We explored several options, including keeping the biologically equivalent dose (BED) equal to that of the standard schedule (Fig 3. B-G), and keeping the total dose a patient receives equal to that of the standard schedule (60 Gy, Fig 3. H,I) as done previously (Leder et al. 2014, Randles et al., 2021). Moreover, we restricted the maximum dose per day to 6 Gy (Wang et al., 2015) and the total treatment duration to 6 weeks.

When investigating determinants of the optimum schedule, we found that the shape of the optimal schedule (i.e. the dose-time curve) sensitively depends on the rate of treatment-independent dissemination. If cell escape in the absence of irradiation is of comparable magnitude as the radiation-induced escape, then de-escalating schedules are optimal, and dose-dependent changes in cell escape have little effect on the optimal schedule (see similarity of schedules in Fig. 3B, C, E). In settings in which cell escape in the absence of irradiation is negligible relative to the radiation-induced escape, whether the optimal schedule changes from a de-escalating to an escalating pattern depends on the manner in which cell escape increases with the radiation dose (Fig. 3D, F,H,I). SI2 provides a detailed characterization of optimal schedules.

### Optimal Schedules for NSCLC treated with radiation

We then used the computational modeling framework to identify optimal fractionation schedules for NSCLC treatment. The model was parameterized using cell motility data of irradiated A549 cells, an NSCLC line exhibiting a positive correlation between radiation dose and migration speed (Fig. 2B). We identified an optimal 2-week radiation schedule with a total dose constraint of 20 Gy and a 6 Gy maximum daily dose. To address the clinical needs of patient populations with lower levels of normal tissue tolerance to radiation, we also identified suboptimal schedules constraining the maximum daily dose to 3, 4, and 5 Gy, which all follow the de-escalating dosing trend (Fig. 4A). To validate the efficacy of these identified schedules, we compared the post-treatment cell counts of the primary tumor cells and escaped tumor cells, averaged over 256 instances of the stochastic simulations, for each schedule (Fig. 4B). The optimal and suboptimal schedules indeed resulted in lower numbers of post-treatment tumor cells at the primary tumor site and lower total numbers of escaped cells. The evolutionary dynamics of the primary and escaped tumor cell populations simulated by the agent-based model demonstrated qualitatively consistent trends with the cell population dynamics predicted by its ODE specification (Fig. 4C and D). The agent-based simulations of treatment response exhibit variability in cell numbers arising from the stochastic nature of cell migration and response to radiation (Fig. 4C). We conducted statistical analyses to further investigate the improved treatment outcomes of the optimized schedules (SF 7A and B) and found that the optimal schedule led to a significant improvement in both tumor shrinkage (*P* < 10^−5^, two-tailed Mann-Whitney U test) and preventing cell escape in comparison to the standard schedule (*P* < 10^−5^). Similar significantly improved outcomes were observed with the suboptimal schedules.

**Fig. 4.**
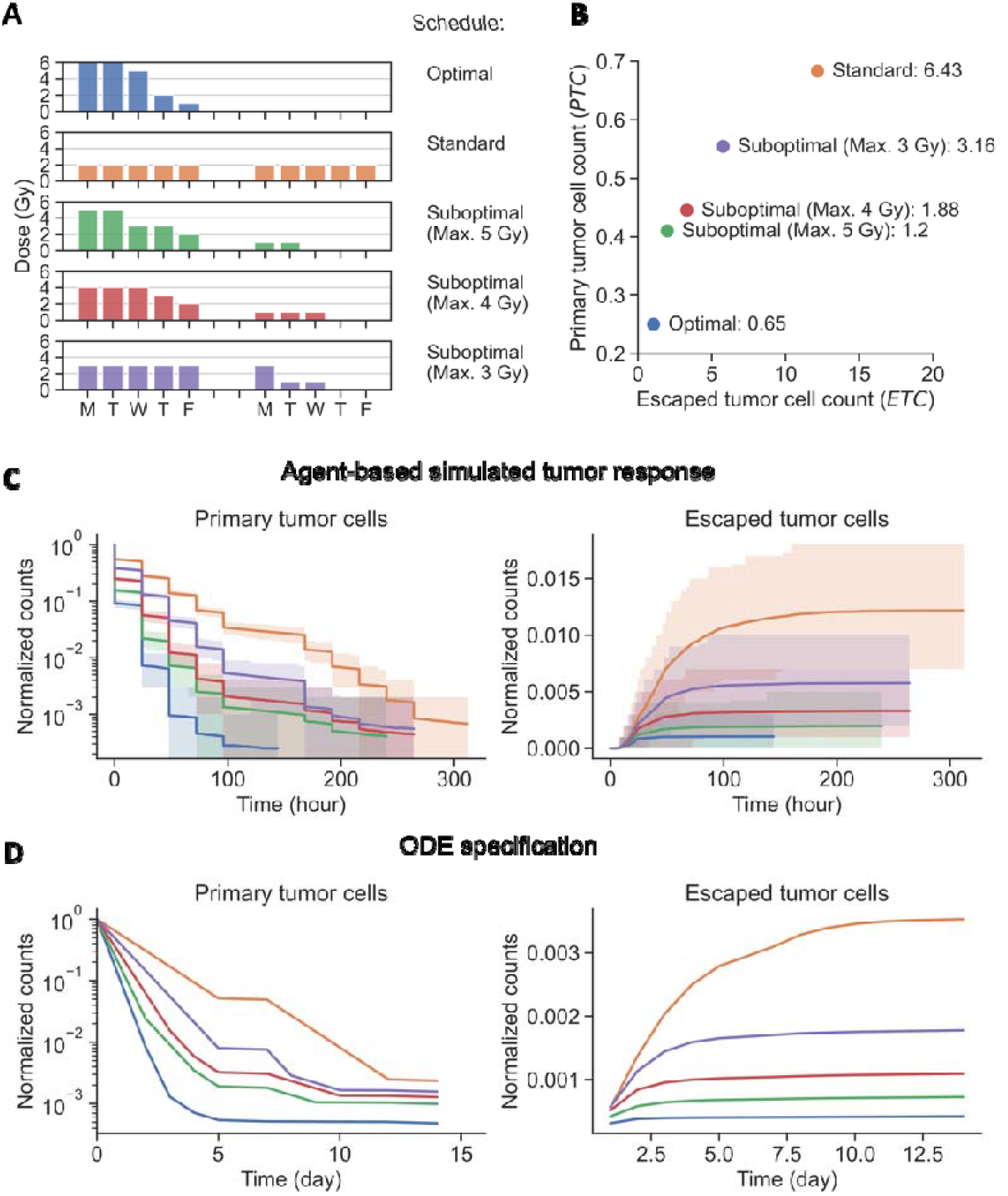
A computational modeling framework identifies optimal and suboptimal radiation administration schedules for NSCLC patients. **(A)** Two-week schedules for validation in simulations. Schedules were optimized given the dose-escape relationship parameterization from the in vitro motility experiments. Different dose-per-day constraints are considered. **(B)** Final absolute numbers of viable cells in the primary tumor and escaped cells after 2 weeks, averaged over 256 simulation runs each initialized with 1000 viable cells. **(C) & (D)** Evolution of the normalized counts of viable primary tumor cells and escaped cells under the radiation schedules shown in panel A, predicted by the spatial agent-based model (shown in **(C)** and its ODE specification (shown in **(D)**) (Methods). The colors of the cell count trajectories correspond to the colors of the radiation schedules in panel A. Shaded regions in **(C)** identify the 5th to 95th percentile ranges and the curves correspond to the mean trajectory for each schedule.

We then performed a local sensitivity analysis to examine the impact of different model parameters on the predicted numbers of primary tumor cells and escaped cells (Methods, SI3). We found that the radiosensitivity parameters *α* and *β* had the largest impact on tumor shrinkage (SF 7C). Furthermore, post-treatment primary tumor cell counts were sensitive to *α* and *β* and insensitive to the ratio of IR margin to tumor radius and the cell motility parameters *a* and *b*. Finally, the number of escaped tumor cells was sensitive to all parameters tested in the analysis, with the baseline speed *a* being most influential.

### *In Silico* Clinical Trial

We then conducted an *in silico* clinical trial with 1,000 simulated patients to further validate the efficacy improvement of our optimized schedules and assess the robustness of the framework relative to inter-patient variability (Fig. 5A, Methods). For each patient, we sampled parameters of tumor characteristics and treatment response from log-normal distributions centered around parameter estimates for the A549 cell line (Fig. 5B, Methods). We then used the agent-based stochastic model to predict the numbers of primary tumor cells and escaped tumor cells in response to radiotherapy for each of three treatment strategies: the model-identified optimal schedule, the suboptimal schedule with a maximum daily dose of 4 Gy, and the standard schedule. The *in silico* tumor response data showcases the variation in cell populations due to different treatment schedules as well as inter-patient variability and intrinsic stochasticity of tumor response to radiation (Fig. 5C).

**Fig. 5.**
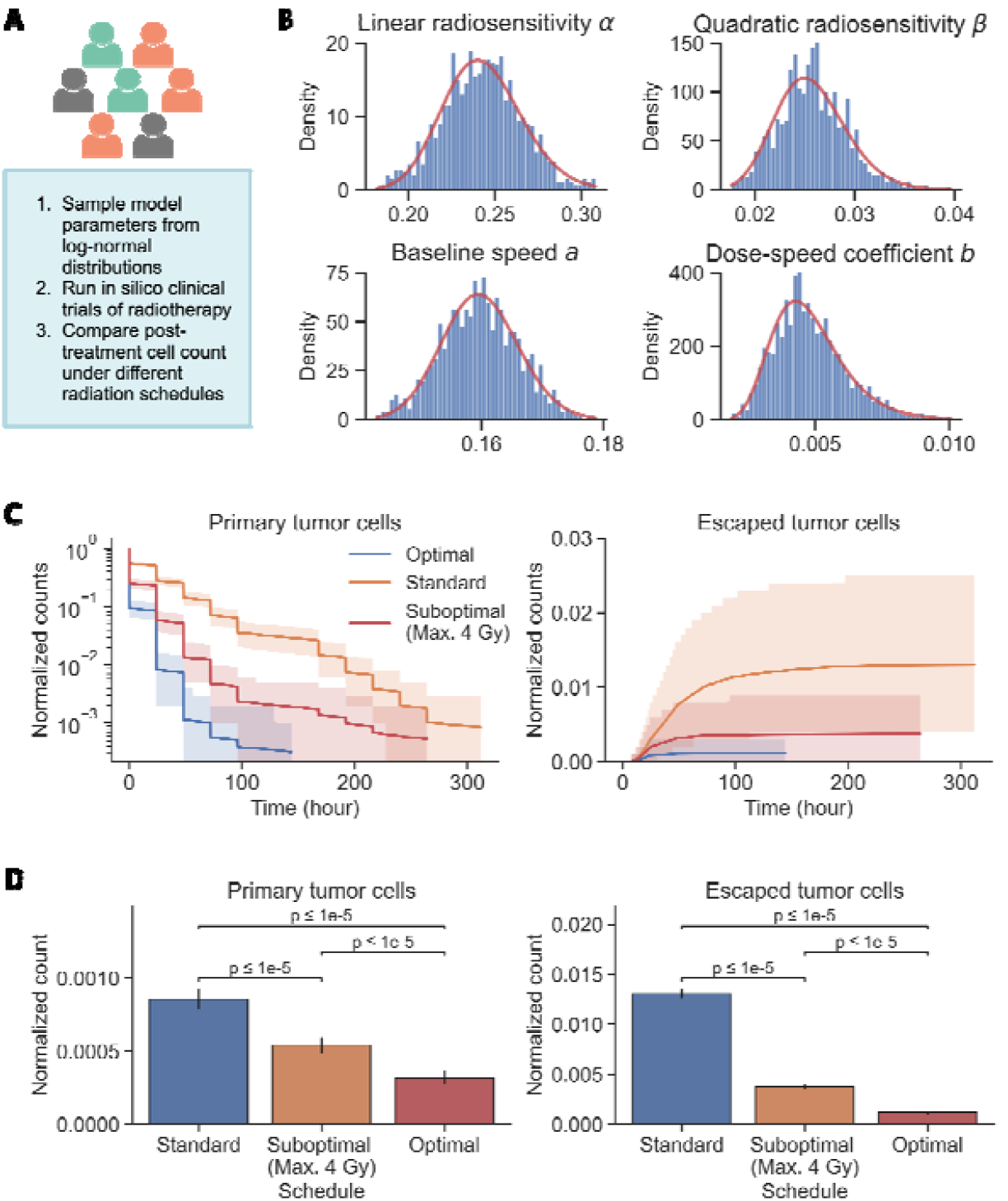
In silico clinical trial of radiation schedules in NSLCL patients. **(A)** Summary of the in silico clinical trial workflow. Colors represent in silico patients with heterogeneous tumor characteristics. **(B)** Histograms of parameter values of linear and quadratic radiosensitivities α and β, baseline speed a, and dose-speed coefficient b used to simulate 1000 in silico patients, along with the log-normal probability density distributions (red curves) used for sampling. **(C)** Cell count trajectories under different fractionation schedules as in Fig. 4A for a simulated patient cohort with varying patient characteristics (see Methods section). **(D)** Comparison of normalized counts of cells in the primary tumor and escaped cells across different schedules from panel A.

We compared the post-treatment cell numbers across the three radiation schedules to assess their effects on tumor shrinkage and escape prevention (Figs. 5D and E). The*−in silico* clinical trial outcomes suggest that both the optimal and suboptimal schedules performed significantly better than the standard schedule at reducing primary site and escaped tumor cell numbers (*P* < 10^−5^, two-tailed Mann-Whitney U test), with the optimal schedule showing a significant improvement over the suboptimal schedule (*P* < 10^−5^). Compared to simulated outcomes under the standard schedule, the optimal schedule resulted in a 62% decrease in primary tumor cell number, averaged over 1,000 *in silico* cases, and a 91% reduction in the number of escaped cells. The suboptimal schedule also showed a substantial efficacy improvement over the standard schedule, with a 37% reduction in primary tumor cell number and a 71% reduction in the escaped cell number. This observation suggests that the novel radiation fractionation schedules predicted by the mathematical framework are considerably more effective in both shrinking primary tumor volumes and lowering the risk of metastatic seeding, while also demonstrating robustness to inter-patient variability and the stochastic nature of tumor cell migration and radiation response.

### Mechanistic machine learning enables parameter identification for personalized schedule prediction

We then sought to investigate whether the features that determine optimal fractionation schedules can be inferred from cell count trajectories over the course of treatment. To this end, we trained a state-of-the-art pretrained transformer model (Hollmann et al., 2025), as well as two baseline machine learning (ML) models for comparison (sparse ridge linear regression and random forest (Breiman, 2001)), on data from our in silico clinical trial (Fig 6A). Specifically, we used the trajectories of cell counts in the primary tumor under the optimal and 4-Gy suboptimal schedules (Fig. 5) to infer the radiosensitivity parameters (*α, β*); these schedules were chosen as they have sufficient variability in administered dose to enable us to disentangle these two parameters. Moreover, we used the trajectories of the number of escaped cells under the baseline schedule, i.e. the schedule for which most cell escape was observed (Fig 5C), to infer cell speed. Across all parameter and schedule combinations, the transformer showed the best five-fold cross-validated test performance (Fig 6B). Notably, the pretrained transformer achieved the best test performance while maintaining similar train and test distributions (Fig 6C), an uncommon outcome in most ML approaches (SF8), suggesting strong out-of-distribution generalizability. This approach enabled us to recover mechanistic parameters with high accuracy, demonstrating that the features which determine optimal radiotherapy schedules can be inferred from longitudinal tumor volume data over short time horizons. Since mapping in vitro characterizations of migratory responses to dynamics in patients remains challenging, this mechanistic machine learning-based approach constitutes an alternative route to clinical application of personalized optimum radiotherapy schedules (Fig 6D).

**Fig. 6.**
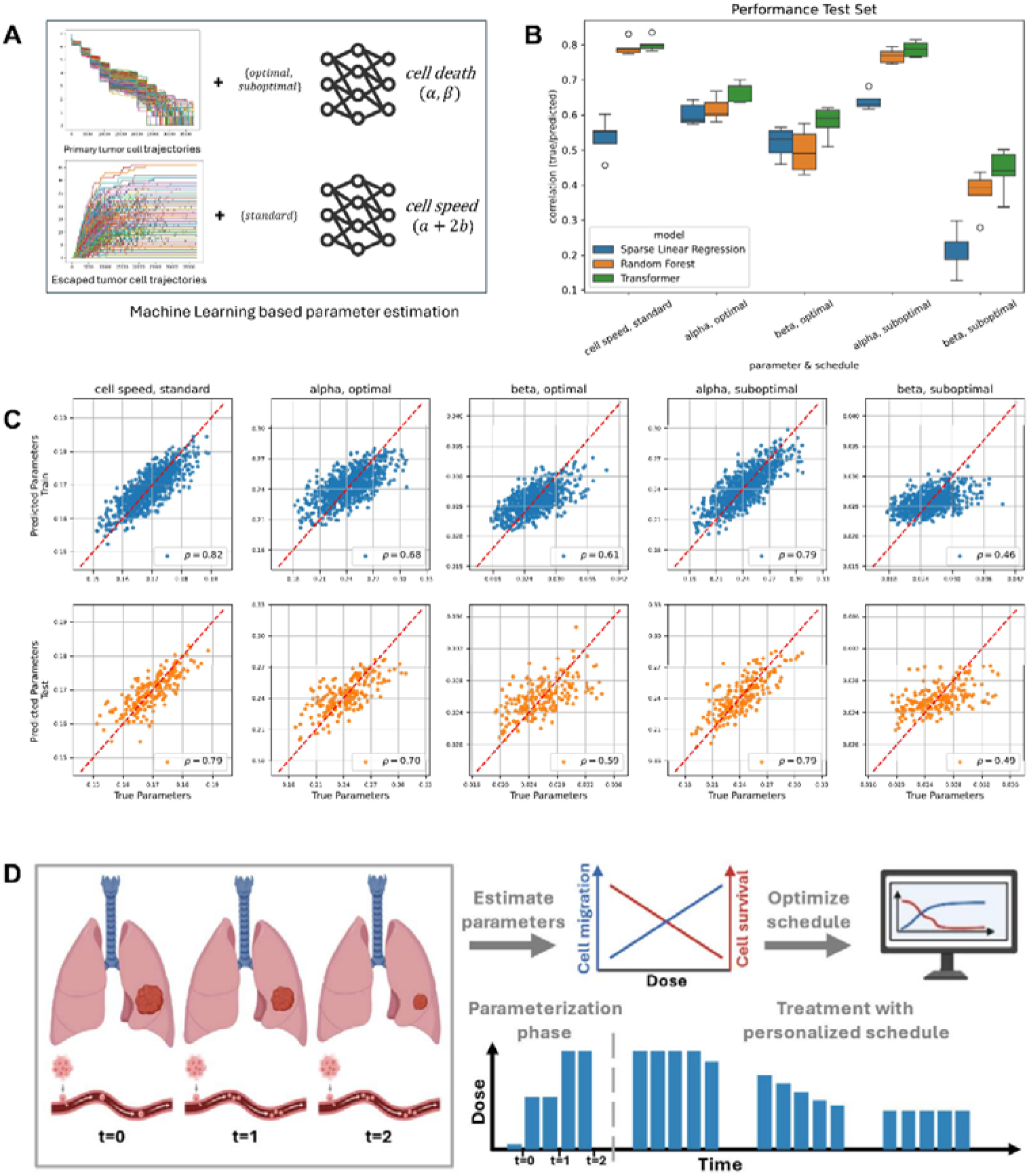
Mechanistic machine learning enables patient-specific parameter estimation. **(A)** Schematic of the inference workflow. As input data, we used cell count trajectories in the primary tumor under the optimal and 4Gy-suboptimal schedule (Fig 5) or cell-escape trajectories under the standard schedule. We then trained different machine learning/artificial intelligence models (Methods) to recover the radiosensitivity parameters or the cell speed (which at a dose of 2 Gy in the standard schedule corresponds to *a* + 2*b*), respectively. **(B)** Five-fold cross-validated test performance, measured as the Pearson correlation coefficient *ρ* between the true parameters and the predicted values (y-axis), across various parameter and schedule combinations (x-axis). Each column corresponds to a unique combination of schedule and inferred parameter. For each setting, we trained three different ML models - Sparse Linear Regression (blue), Random Forest (orange), and a Transformer (green). Each of the models was trained five times on different training and testing splits, with error bars indicating the variability. The Transformer consistently outperformed the other models across all settings. **(C)** Detailed predictive performance for the transformer model, where each column represents different parameter and schedule combinations. The top row shows results for the 800 training trajectories (blue dots), while the bottom row presents predictions for the 200 unseen test trajectories (orange dots). Each scatter plot shows the relationship between true (x-axis) and predicted parameter values (y-axis), where the diagonal red dashed line indicates perfect prediction. The training (top row) and test distribution (bottom row) for the transformer model are very similar in each setting, suggesting strong prediction power and out-of-distribution generalizability for unseen trajectories. **(D)** Envisioned application: optimizing personalized radiotherapy schedules with our machine learning approach. The patient’s tumor volume and CTCs would be monitored during a parameterization phase, to infer migration and radiosensitivity parameters. Based on these parameters, an optimal fractionation schedule would be identified to optimize subsequent fractions.

## Discussion

Radiotherapy is an effective mainstay option for the treatment of most cancer types, but an increasing volume of literature suggests that it may inadvertently increase tumor invasiveness and the risk of metastasis formation by enhancing cell motility. Here, we investigated whether radiotherapy schedules can be optimized to mitigate this therapy-induced risk. We quantitatively characterized the relationship between radiation dose and induced changes in tumor cell motility using live-cell imaging experiments and developed a computational framework to model the dynamics of cell migration in response to radiotherapy. We used this framework to identify optimal treatment schedules and to perform an in silico clinical trial to quantify how optimized schedules improve therapeutic outcomes relative to standard of care radiotherapy schedules in NSCLC. We also demonstrated that we can recover patient-specific parameters used in our in silico clinical trial using mechanistic machine learning, paving the way for potential future clinical applications of this methodology.

In our live-cell imaging experiments, we observed a large extent heterogeneity even between apparently similar cell lines of the same cancer type. For instance, the lung cancer cell lines A549 and NCI-H460 showed opposite migratory responses, despite both harboring KRAS activating mutations (G12S and Q61H, respectively) and loss of CDKN2A and STK11. This heterogeneity is consistent with the variable findings that have been reported in previous studies exploring migration responses in fewer cell lines (Goetze et al. (2007); Kawamoto et al. (2012); Merrick et al. (2021); Ogata et al. (2005); Pickhard et al. (2011); Steinle et al. (2011); Young and Bennewith (2017); Moncharmont et al. (2014)). Moreover, our findings align qualitatively with previous studies finding decreasing motility for MCF7 (Young and Bennewith (2017)) and HCT116 (Goetze et al.(2007)) and increasing motility for A549 (Jung et al. (2007); Tahmasebi-Birgani et al. (2019)).

While ablative regimens are preferred for effective tumor shrinkage, lung tumors in anatomically challenging locations, e.g., abutting critical organs-at-risk such as the heart or central airways, or more extensive in distribution, need to be treated with sub-ablative treatment regimens involving longer courses of daily treatment with smaller fraction sizes in the 2–6 Gy range (Said et al. (2024)). However, rather than keeping doses constant throughout treatment, as is current standard of care, our modeling platform suggests that the optimal fractionation schedule starts with higher daily doses that decrease monotonically over the course of treatment. Compared to the *in silico* outcomes under the standard schedule, the model-identified optimal schedule led to a 62% decrease in the post-treatment primary tumor volume and a 91% reduction in the total number of escaped cells.

In light of the striking heterogeneity in migratory response to radiation both in the literature and in our experimental data, identifying patient-specific migratory responses remains a key challenge on route to clinical implementation of the proposed optimized schedules. Efforts such as live-cell imaging to catalogue migratory response across cancer types, or to identify markers for migratory behaviors, may address this challenge. Yet, in the absence of a reliable catalogue or established biomarkers, an alternative approach to addressing this challenge is to infer such parameters from changes in clinical measurements at the beginning of radiotherapy to enable personalized adaptation of radiotherapy schedules early during treatment (Fig 6D).

Our mechanistic approach modeling combined with ML-based parameter estimation provides a proof of concept that such inference is possible based on raw trajectories of cell counts in the primary tumor and of escaped cells. Already with small numbers of escaped cells and trained on few trajectories, our mechanistic and data-driven pipeline identifies nuanced differences in the patient-specific parameters over a two-week treatment horizon. This characteristic makes our integrative predictive platform attractive for future clinical implementation of personalized schedules.

Our findings have several caveats, including the use of in vitro measurements of cell motility in a sparsely populated dish to parameterize our computational framework. This approach assumes that these findings mimic changes in motility in densely populated tissues, and more generally that these changes in motility are the mechanism by which radiation leads to an increased risk of metastatic seeding. While other mechanisms (Moncharmont et al., 2014), such as changes in cell adhesion or interactions with the extra-cellular matrix, could be integrated into our framework, quantitatively measuring these mechanistic effects would require further experiments. Furthermore, our experimental data only provides estimates for motility changes following a single dose of radiation. We therefore investigated schedules with at most one dose per 24 hours (the timeframe of our experiment), and considered motility impacts of a given dose to be independent of the previous doses of radiation that a tumor had received. Further experiments are needed to determine multi-dose radiation response.

In summary, we created a predictive computational modeling platform to analyze tumor cell numbers and their spread during radiotherapy. We identified an optimal treatment schedule using data from the lung cancer cell line A549 and conducted extensive *in silico* clinical trials, incorporating variations in radiation response and other tumor characteristics. By training transformer models on this *in silico* data, we were able to accurately recover mechanistic parameters, showing that the key factors influencing optimal radiotherapy can be deduced from longitudinal tumor data. This integrative predictive approach provides a foundation for rationally designing optimal clinical trials across different cancer types.

## Methods

### Cell culture experiments

Parental cell lines (A498, A549, NCI-H460, HCT116, LOX-IMV, MALME-3E, MCF7, SK-MEL5, U2OS, UACC257, UACC62, and UO31) were obtained from the American Type Culture Collection. The cells were thawed and propagated in RPMI (Gibco) with 5% fetal bovine serum (FBS). A full description of the cell culture experiment is available in Stewart-Ornstein and Lahav (2017).

### Measurement of cell motility following irradiation

Cells from the 12 cell lines were exposed to radiation with doses 1, 2, 4, 6, and 8 Gy and imaged every 15 minutes for 24 hours post-irradiation using live-cell fluorescence microscopy, which was performed following the procedure outlined in Stewart-Ornstein and Lahav (2017). To track the motion of individual cells, live-cell imaging data were processed using a custom MATLAB code, where the phase images were used to manually identify and track cells over time. Then, the 2D coordinates of cell centroids were extracted at each time point. Cells that died or moved out of the field of view during the experiment were excluded from our analysis. The single-cell trajectories were constructed by linearly interpolating between cell locations at consecutive time points. Detailed methodology for irradiation, live-cell microscopy, single-cell tracking, and live-cell measurement of p53 dynamics can be found in Stewart-Ornstein and Lahav (2017) and Reyes et al. (2018).

### Statistical analyses of cell motility and p53 dynamics

The average speeds of individual cells were estimated from their single-cell trajectories, which in turn were obtained from live-cell imaging and manual segmentation. For each cell line, we calculated the Pearson correlation coefficient between radiation dose and average cell speed along with the associated p-value; a least squared linear regression of dose against speed was also performed. The statistical analyses were conducted in R v.4.4.1. *P <* 0.05 was considered statistically significant in all statistical tests.

### Computational modeling

To examine the effects of radiotherapy on tumor shrinkage and cell migration, we developed a stochastic agent-based model of radiation treatment response based on a spatially explicit computational model we previously developed for glioblastoma treatment response modeling (Randles et al., 2021). In our model, we considered a 2D cross-sectional area of a spherical tumor, where the circular cross-section consists of a population of 1,000 non-overlapping tumor cells. These tumor cells are constantly in motion due to both cell-cell interactions and random Brownian motion. Following exposure to ionizing radiation, the irradiated tumor cells either enter quiescence with halted proliferation or become apoptotic due to DNA damage. The tumor cells surviving the irradiation migrate with dose-dependent changes in cell speeds.

### Radiotherapy

In this model, radiation-induced cell death is represented by the linear quadratic model (Hall and Giaccia, 2012, Fowler, 2010, and McMahon, 2018). According to this model, following irradiation with dose *d*, cell death occurs with a probability

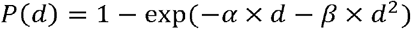

where *α* and *β* are the respective linear and quadratic radiosensitivity parameters. The dying cells lose cellular functions including growth, division, and aging. The remaining tumor cells that have survived irradiation enter a quiescent state, where they undergo cell cycle arrest to repair DNA damage. After some time, surviving tumor cells may re-enter the cell cycle and resume proliferation (Fig. 2A). The duration of the cellular arrest and repair in cancer cells can vary, typically ranging from days to weeks. Therefore, it is reasonable to assume in our model that, once irradiated, tumor cells remain in a quiescent state throughout the course of radiotherapy, given that the interval between radiation administration times do not exceed two days.

### Cell migration

The movement of tumor cells is driven by random Brownian motion and cell-cell interactions. At each time step, the positions of individual tumor cells are updated based on random displacements in *x* and *y* directions. The displacements are sampled from a normal distribution, i.e., 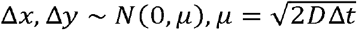, where Δ*x* and Δ*y* denote the respective cell displacements in *x* and *y* directions, _Δ_*t* denotes the simulation time step size, and *D* is the diffusion coefficient of the random Brownian motion. The speed of random movement of tumor cells is governed by the step length parameter *u*. Prior to radiation treatment, we prescribe a baseline speed a for the simulation of Brownian motion. To account for the dose-dependent migratory behavior of tumor cells, we assume that following irradiation of dose *d*, the surviving tumor cells (normal or quiescent) migrate in Brownian motion with step length *u* = *a* + *b* × *d*, while the radiation-killed cells migrate in Brownian motion with baseline speed *u* = *a*. Additionally, each pair of tumor cells in the model undergoes either attractive or repulsive interaction forces derived from the Lennard-Jones potential, which is a function of cell sizes and distance between the centers of cells. The interaction forces, consisting of short-ranged repulsion forces and long-ranged attraction forces, move tumor cells accordingly. The repulsion forces ensure that tumor cells do not physically overlap in the 2D domain, while the attraction forces simulate the adhesion between interacting tumor cells (Verlet, 1967, Randles et al., 2021). The cell migration driven by the cell-cell interactions is unaffected by irradiation in the model. A primary tumor region is defined based on the pre-treatment tumor size with a 10% margin extending outside of the tumor boundary. During a treatment response simulation, the tumor cells that migrate outside of the primary tumor region are considered escaped and removed from the system.

### Quantifying treatment objectives

To incorporate the dual treatment objectives of reducing tumor sizes and minimizing metastasis risk into our framework, we defined a score function *V* based on the post-treatment tumor cell number and the total number of cells that have escaped the primary tumor region, i.e., *V* = (1− *ω*) × *PTC* + *ω* × *ETC*, where *PTC* denotes the number of tumor cells in the primary tumor region at the final simulation time, *ETC* the total number of escaped tumor cells, and _*ω*_ is a weight parameter that describes the relative importance of tumor shrinkage and metastasis prevention in setting treatment goals. Given a candidate radiation fractionation schedule, a score is calculated using the value function, providing a quantitative evaluation of the schedule’s treatment performance.

The results presented throughout were determined with an equal weighting of cells in the primary tumor region and escaped cells (*ω*= 0.5). Given a fixed BED, decreasing *ω* reduces the extent of the shift towards (de-)escalation, so that the schedule becomes more similar to the uniform standard of care schedule, while increasing w increases the shift towards (de-)escalation. To address the stochasticity in the agent-based simulations, we performed 256 instances of each schedule and calculated scores using the averaged results for model validation and sensitivity analysis (Figs. 4B-D and SF7).

### Optimization

For computati<u>o</u>nal efficiency, we identify optimal treatment schedules in a simplified ODE specification of our model. This specification tracks cell movements between *N* compartments, initialized with mas<u>se</u>s 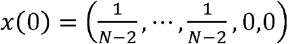, where the first *N*−2 compartments represent the initial tumor region, with different layers being at different proximity to tumor margins. The second-to-last compartment represents the IR margin, and the last compartment the escaped state. Doses are administered at discrete time points spaced in one-day intervals and shrink the tumor mass in the first *N*−1 compartments according to the linear quadratic model. Between doses cells proliferate at a rate *g*, and unless they have escaped, migrate between neighboring compartments at a rate *k*_*m*_, so that we obtain a linear first order system with constant coefficients.

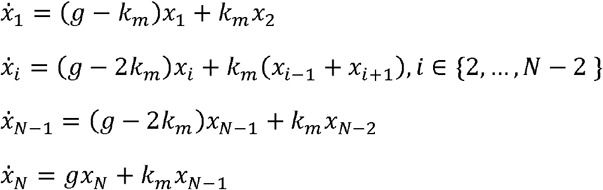

This system has an explicit solution in terms of its matrix of coefficients, and by concatenating the tumor shrinkage and the subsequent growth and movement at each irradiation time point, we can efficiently compute the trajectory of the tumor mass distribution for a given schedule. The migration or escape rate *k*_*m*_ is a function of the most recent administered dose. Doses on weekends are set to zero. The doses at each remaining time point are then optimized using the Trust-Region Constrained Algorithm (TRCA) implemented in the ‘scipy.minimize()’ function in SciPy version 1.14.0, with daily doses bounded between 0 Gy and 6 Gy, and with an equality constraint ensuring that either the BED or the total cumulative dose is kept constant. Given that TRCA is a local optimization algorithm, our objective is to identify schedules that are approximately optimal and significantly outperform the standard schedule. The schedules in Fig. 3 correspond to parameters *N* = 6, *g* = 0 and *k*_*m*_ as the escape rate plotted above each schedule. The dose-escape curve in Fig. 3E corresponds to curve that we estimate from our experimental data. This curve (*k*_*m*_) is also used in Fig. 4A-B, along with the same parameters *N* = 6 and *g* = 0. For clinical applicability, doses in 4A are rounded to integer Gray values.

For a given weight *ω* on escaped cells in the value function, increasing the parameter *N* increases the number of cells that are at little risk of escape during the course of treatment and therefore flattens the optimal treatment schedule. Increasing *ω* counteracts this effect sufficiently much so that for *ω* close to 1, the optimal schedules in Figs. 3 and 4 are not sensitive to *N*.

Setting *g* to 0 matches the common clinical observations that viable cells that have recently been exposed to radiation tend to be in a senescent state – which we also mimic in the full agent-based model. Generally, the optimal schedules tend to only be sensitive to *g* in extreme cases where the schedule at *g* = 0 puts the dose on some days very close to zero (these cases arise at small *N* and if *k*_*m*_ is a very steep function of dose), so that at *g* > 0 the tumor shrinkage brought about by the dose is smaller than the subsequent re-growth of the tumor. In those cases, increasing *g* thus flattens the optimal schedule somewhat, so as to prevent such net tumor growth.

### Model parameterization

The implementation of the Brownian motion of cells in the agent-based simulations was parameterized by the time course motility measurements of 301 A549 cells. We calculated the mean squared distance (*MSD*) of A549 cells irradiated with doses ranging from 1 Gy to 8 Gy from the positions of cell centroids measured every 15 minutes for 24 hours. Then, we estimated the Brownian diffusion coefficient *D* of cells treated at each dose level using the formula *D* = *MSD*/(4*t*) and regressed dose level against the step length 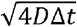. The resulting dose-dependent step length is given by 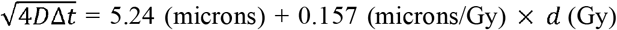, which was implemented in the agent-based model to simulate the dose-dependent cell migration. Note that since we do not have the cell motility measurements of un-irradiated A549 cells, for baseline Brownian motion, we used the estimated step length (5.24 microns) at 0 Gy from regression. The values of radiosensitivity parameters *α* and *β* were obtained by fitting the linear-quadratic model to the clonogenic survival data of A549 cells (Bromley et al., 2009).

### Sensitivity analysis

The model sensitivity analysis was conducted using a derivative-based method, where we varied the parameters values of the baseline speed *a*, dose-speed coefficient *b*, linear and quadratic radiosensitivities *α* and *β*, and the IR margin ratio by ±10%, ±20%, ±30%, and ±40%. Then, we simulated the tumor response to the optimal radiation schedule 256 times for each perturbation level. The sensitivity of model parameters was evaluated by approximating and averaging their first-order partial derivatives. Specifically, for each model parameter, we calculated their MSIs to the averaged and normalized counts of primary tumor cells (*PTCs*) and escaped tumor cells (*ETCs*). The formula of *MSl*s is given by

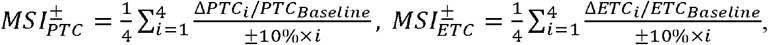

where *PTC*_*Baseline*_ and *ETC*_*Baseline*_ denote the model outputs (normalized cell counts averaged over 256 simulation runs) with unperturbed parameters, and Δ*PTC* and Δ*ETC* denote the changes in model outputs due to the ±10% ×, perturbation of a parameter. Further, we examined the importance of each parameter by performing univariate linear regressions of model outputs on parameter values at different perturbation levels. Parameters that are statistically significant at the 5% level are defined to be sensitive.

### *In silico* clinical trial

The *in silico* clinical trial was conducted with 1,000 simulated patients, where we defined each patient with a unique set of parameters (baseline speed *a*, dose-speed coefficient *b*, and radiosensitivity parameters *α* and *β*) sampled from log-normal distributions (Fig. 5B) informed by experimental data (Bromley et al., 2009) and cell motility estimation. We compared the tumor response in simulated patients treated with the optimal schedule, the suboptimal schedule with a maximum daily dose of 4 Gy, and the standard schedule. Two-tailed Mann-Whitney U tests were used to evaluate differences in post-treatment cell numbers and escaped cell numbers.

### Machine learning-based parameter estimation

We considered the cell count trajectories 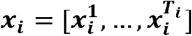 and the escalated cell count trajectories 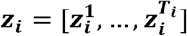 for each patient *i* ∈ 1,…, *N* with observation length ***T***_***i***_.. Each of the simulated *N* = 1000 trajectory combinations per treatment regime ***Q*** was labelled with the corresponding parameters 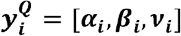.We learned a statistical machine learning (ML) model **ℳ** by regressing the parameters 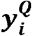 onto the cell trajectories ***x***_***i***_ and ***z***_***i***_; that is, we learned three models 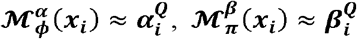, and 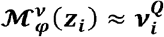, for each of the parameters ***α, β, v*** and each treatment schedule ***Q***, as depicted in Fig. 6A. Each ML model 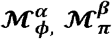, and 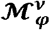, was parametrized by some parameter sets (***ϕ, π, v***) which were learned by 3 different ML algorithms: sparse ridge linear regression, random forest (Breiman, 2001), and pretrained transformers (Vaswani, 2017; Hollmann et al., 2025). For model training, we split the *N*=1000 patients into 5-folds of 200 patients each. We then trained each modedl on the 800 patients and evaluated the performance of the remaining 200 test patients, and repeated that approach 5 times. Hyperparameters of the ML algorithms were slightly tuned in the inner loop of nested cross-validation. We used the sklearn (Pedregosa, 2011) package in python sparse linear regression and the pretrained transfer from the package tabPFN (Hollmann et al., 2025). The full code and the trained models with the exact hyperparameters and implementation details are provided for reproducing the ML parameter estimation experiments. To measure the performance, we used as metric the correlation 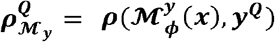 for each parameter ***α, β, v***, for each ML model under consideration (sparse ridge linear regression, random forest, and pretrained transformers), and for the train and test data, respectively.

## Declarations

D.K. is a consultant for AstraZeneca and Genentech/Roche. D.K. declares that none of these relationships are directly or indirectly related to the content of this manuscript.

F.M. is a co-founder of and has equity in Harbinger Health, has equity in Zephyr AI, and serves as a consultant for both companies. She is also on the board of directors of Recursion Pharmaceuticals. F.M. declares that none of these relationships are directly or indirectly related to the content of this manuscript.

## Code Availability

All code for this study is available under https://github.com/Michorlab/radiotherapy-induced-cell-migration.

