## Supplementary Info for "Mechanistic modeling and machine learning identifies optimum radiotherapy schedules to prevent treatment-induced metastasis"

Supplementary information

**1 Cell migration patterns and p53 dynamics following irradiation**

We examined the heterogeneity of cell migration patterns following irradiation, observing that the migration speeds of irradiated cells remain predominantly homogeneous over time throughout the 24-hour experiment (Fig. S1). This finding indicates a time-consistent effect of radiation on cell migration, rather than an initial surge in motility due to DNA damage which is mostly repaired within 24 hours (McMahon et al. (2016)). Furthermore, we found that the variability in cell motion following irradiation remains similar over time across different cell lines and doses (Fig. S2). Our examination of the most migratory cells in each cell line demonstrated consistent dose-speed relationships compared to the general population, i.e., for cell lines with non-significant or negative dose-speed correlations, no significant increase in speed with increasing dose was detected among their fastest moving cells (Fig. S3). Cell dispersal from their original locations varied substantially across cell lines and doses, yet no directional preference was observed (Fig. S4). This finding suggests a lack of collective motion on a population scale.

Additionally, we performed linear regressions of cumulative p53 level against dose and speed against cumulative p53 level and calculated Pearson correlation coefficients (Fig. S5 and 6). Notably, the UO31 and A549 cell lines that showed significant positive dose-speed correlations also exhibited significant positive correlations between cumulative p53 level and dose as well as cell speed and cumulative p53 level. To further investigate the role of p53 in radiation-induced increases in cell motility, we performed a linear regression of speed against dose and the interaction between dose and cumulative p53 level for the UO31 and A549 cell lines (Tab. S1). The dose-p53 interaction had a significant positive effect on speed in A549 cells (*P* = 0.048), while no significant effect of the interaction was detected in UO31 cells (*P* = 0.15).

In our analyses, cumulative p53 levels were obtained by summing the p53 levels across all time points. Detailed methodology for live-cell measurement of p53 dynamics can be found in Stewart-Ornstein and Lahav (2017).

**2 Characterization of optimal treatment schedules**

If radiation leads to an increased risk that cells disseminate and seed metastases, then there may be a tradeoff between shrinking the tumor effectively and containing the risk of induced metastases. However, the tradeoff, and whether it exists, depends on details of these dynamics and their interactions. For instance, while irradiating with lower doses may initially induce less cell escape from the primary tumor, it also leaves more viable tumor cells that are at risk of induced metastatic seeding later on during the treatment. Here, we provide a thorough characterization of the conditions leading to a trade-off between induced metastasis risk and tumor shrinkage so that lower doses at the beginning of treatment (escalating schedules) are optimal, versus when both tumor shrinkage and long-term cell escape are minimized by high doses at the beginning of treatment (de-escalating schedules). To this end, we consider optimal schedules under specific functional forms for the dose-escape relationship as presented in the main text (Figure 3). The key feature determining whether optimal schedules are de-escalating or escalating is whether cell escape increases disproportionately with the administered dose. To provide mathematical intuition for this result, we designed a much simplified but analytically tractable version of our model.

In the main text we distinguish between constraining either the total BED or the total cumulative dose that a patient receives. Let us here first consider a scenario in which the BED a patient receives is constant and let us assume that the risk of metastatic seeding is negligible. In this scenario, the decision which schedule optimally targets the primary tumor depends on the ratio of the coefficients of the first and second order terms in the linear quadratic model ($\alpha/\beta$ ratios) of the tumor and of the surrounding tissue. Concretely, if the $\alpha/\beta$ ratio of the tumor is lower than that of the surrounding tissue, a hypofractionated schedule is optimal (Ritter, 2008), consisting of the maximum tolerable dose per day over a short time period. Conversely, if the $\alpha/\beta$ratio of the surrounding tissue is lower than that of the tumor, the optimal schedule is hyperfractionated. The latter is generally observed for NSCLC, with $\alpha/\beta$ratios around 9.5 measured in A549 cells (Bromley et al., 2009) and around 3.3 for normal lung tissue (Van Dyk et al., 1989), so that the uniform, low-dose (2Gy) standard of care schedule performs best.

However, if the risk of cells escaping the primary tumor over the course of treatment is not negligible, our model shows that the standard uniform schedule is no longer optimal and that the total number of surviving tumor cells is lowest under schedules that initiate with higher fractional doses (Fig. 3B). This result does not require that cell escape varies with the administered dose. Intuitively, this shift to more aggressive dosing at earlier time points becomes optimal because delaying doses comes at the cost of not being able to target cells that have escaped the irradiated region before receiving radiation.

In situations in which cell escape is non-negligible in the absence of radiation but increases with the administered dose (Fig. 3C), the optimal schedule retains this de-escalating shape. However, if cell escape in the absence of radiation is negligible and increases in the same manner with the administered dose, then the optimal schedules are escalating (Fig. 3D). This strong dependence of the optimal schedule on the extent of cell escape absent radiation arises when the additional cell escape per additional Gy is highest at high doses, i.e. if cell escape increases convexly with the dose (Fig. 3C,D). If the dose dependency does not increase at higher doses, then de-escalating schedules are optimal independent of the extent of dose-independent cell escape from the primary tumor site (Fig. 3E, F). Since we optimized schedules with identical weights on escaped cells and remaining cells in the primary tumor (Methods), all schedules achieve a reduction in the total number of cells (Fig. 3G). This reduction is substantial, reducing the relative cell escape to less than 40%, and the relative total number of remaining cells to less than 25% compared to the standard of care schedule.

If, rather than keeping the total BED consistent, the total absolute dose is constant, we observe a similar pattern for the optimal schedules. In settings with negligible cell escape in the absence of radiation and ever more strongly increased cell escape at the larger doses, escalating schedules are optimal (Fig. 3H). In all other settings with cell escape, de-escalating schedules are optimal. However, this latter shift toward de-escalation does not manifest in a change from the schedule that is optimal in the absence of cell-escape, as this schedule is already maximally hypofractionated (Fig. 3I; constraining total dose is equivalent to setting $\beta$=0 for the normal tissue).

Let us now simplify the framework to focus on how the shape of the dose-escape relationship relates to this tradeoff and the dichotomy between regimes in which de-escalating schedules are optimal vs regimes in which escalating schedules are optimal. Rather than explicitly describing the discrete time points at which doses are administered, we consider radiotherapy schedule to be described as a function in continuous time, and for mathematical simplicity we consider treating the tumor over an infinite time horizon. Moreover, we do not take into account spatial patterns explicitly and instead assume that the rate at which cells escape the tumor only depends on the radiation dose. Since we can express both the death rate and the escape rate as monotone functions of the dose, we can directly express the net growth (or shrinkage) rate γ(*t*) at time *t* as a function of the chosen escape rate *s*(*t*) – at every point in time, we are choosing a point on a growth-speed curve. Lastly, we do not impose any dose-constraints and require only that the targeted tumor mass m(t) goes to zero over the course of treatment, lim_t→∞_ *m*(*t*) = 0. Without a dose constraint the speed of tumor shrinkage is of course maximized by maximally high doses. To identify when maximizing tumor shrinkage and minimizing cell escape conflict, we thus need to identify when cell escape is *not* minimized by maximally high doses.

The total cell escape over the course of treatment is given by the integral over the product of cell mass and escape rate *m*(*t*)*s*(*t*). Using the equation determining how *m*(*t*) evolves in terms of *γ*(*t*) and *s*(*t*) we obtain the following minimization problem.

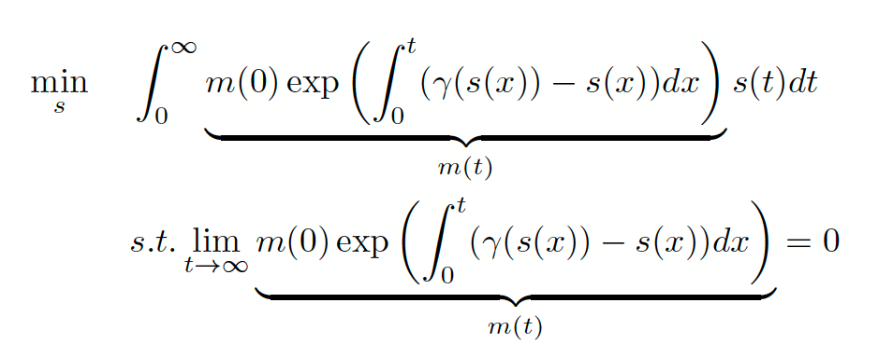

Note that this problem is stationary, so that the function *s*(*t*) that minimizes this expression does not depend on *t*, and not on *m*(0). Under an optimized dose schedule, the integral over *m*(*t*)*s*(*t*) therefore reduces to the following, much simpler expression:

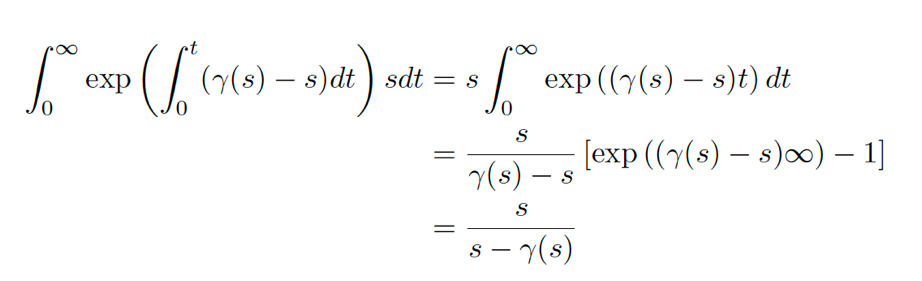

For *s*, only positive values make sense. Moreover, the requirement lim*_t→∞_ m*(*t*) = 0 implies that in the stationary solution we must have *γ*(*s*) *− s <* 0.

As both *γ* and *s* are functions of the administered dose, they exist on some bounded domain determined by biological dose constraints. We can now formulate conditions for minimizers (*γ^∗^*(*s*)*, s^∗^*) of the above integral on the interior of this domain. If an interior solution exists, then the minimal overall cell escape is achieved by balancing the instantaneous effect of radiation on cell escape, with the long-term effect of leaving more viable cells that may escape later during the course of treatment. If no interior solution exists, the latter effect universally dominates the former effect^[1]^.

Assuming differentiable *γ*, a necessary condition for the above integral to be minimal^[2]^ is given by

$$s=\frac{\gamma^{*}(s)}{\gamma^{*'}(s)}$$

Note that *γ*(*s*) and *γ^′^*(*s*) are negative per assumption – we aim to shrink the tumor, and more shrinkage (smaller *γ*) comes at the cost of more movement. This extremum is a minimum if *γ^∗^′′* (*s*) *>* 0, i.e. if *γ* is locally convex.^[3]^

This finding mirrors results in the main text (Fig. 3). For an almost linear dose-escape function, the concavity of the linear-quadratic model makes the escape-survival mapping concave, so that lowering instantaneous (early) doses yields no long-term reduction of escaped cells. If the dose-escape function is strongly convex, so that the escape-survival mapping is convex, smaller early doses become optimal.

**3 Sensitivity analysis**

The parameters examined in the sensitivity analyses include the baseline speed $a$, dose-speed coefficient $b$, linear and quadratic radiosensitivities $\alpha$ and $\beta$, and the IR margin ratio. We found that the post-treatment primary tumor cell number is sensitive to $\alpha$ and $\beta$ and insensitive to the IR margin ratio and the cell motility parameters $a$ and $b$, which is expected since the $\alpha$ and $\beta$ parameters are directly responsible for the cell killing at primary tumor sites. The number of escaped tumor cells is sensitive to all the tested parameters, where the baseline speed $a$ is substantially more influential than the other parameters (Fig 5). This finding implies that the innate cell motility of tumor cells is highly associated with the likelihood of cell escape and metastatic seeding. While the dose-speed coefficient $b$ is a sensitive parameter, its impact on cell escape during radiotherapy is far less than that of the baseline speed a. This observation indicates that the A549 cells are inherently at risk of metastatic seeding, and such risk is relatively unaffected by the radiation doses and dose-dependent cell motility changes. The number of escaped cells is also sensitive to the IR margin ratio, defined as (IR radius/tumor radius -1), as it directly determines how far a tumor cell needs to travel away from the primary tumor mass to be considered escaped. The influences of radiosensitivity parameters $\alpha$ and $\beta$ can be explained by that efficient cell killing due to high radiosensitivity reduces the amount of cells that are viable and able to escape.

**4 Supplementary figures and table**

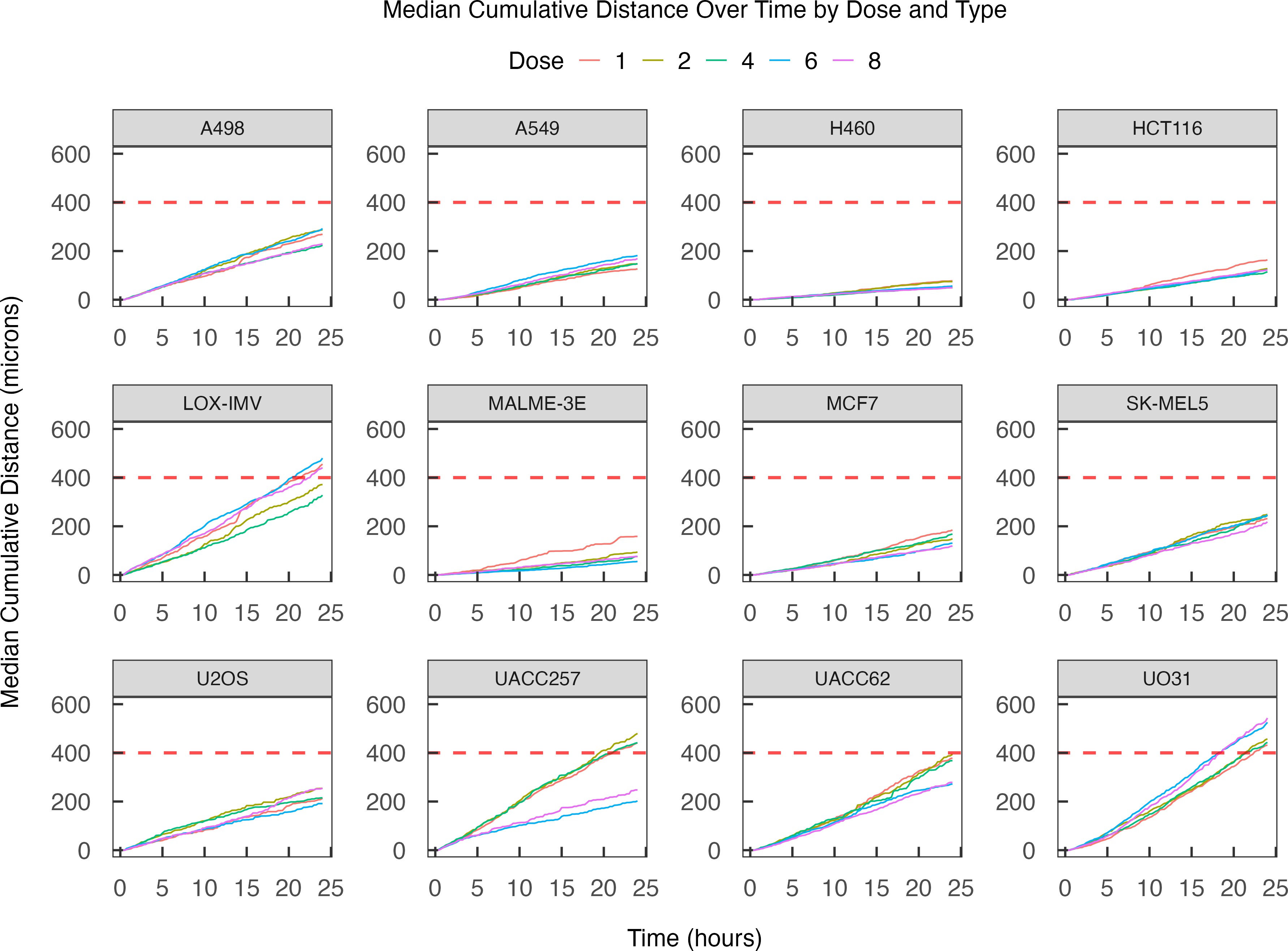

**Supplementary Fig. 1 The median cumulative distance traveled by cells over time, grouped by radiation dose and cell line.** The migration speeds of irradiated cells appear largely constant throughout the 24-hour experiment period, which is reflected in the linear trajectories of cumulative distances. This observation suggests that migration speeds remain temporally homogeneous, rather than exhibiting an initial surge in response to radiation-induced stress followed by a decrease after stimulus removal. Among the 12 cell lines, LOX-IMV, UACC257, and UO31 traveled the longest migration distances following irradiation, with median cumulative distances exceeding 400 microns (indicated by the red dashed line).

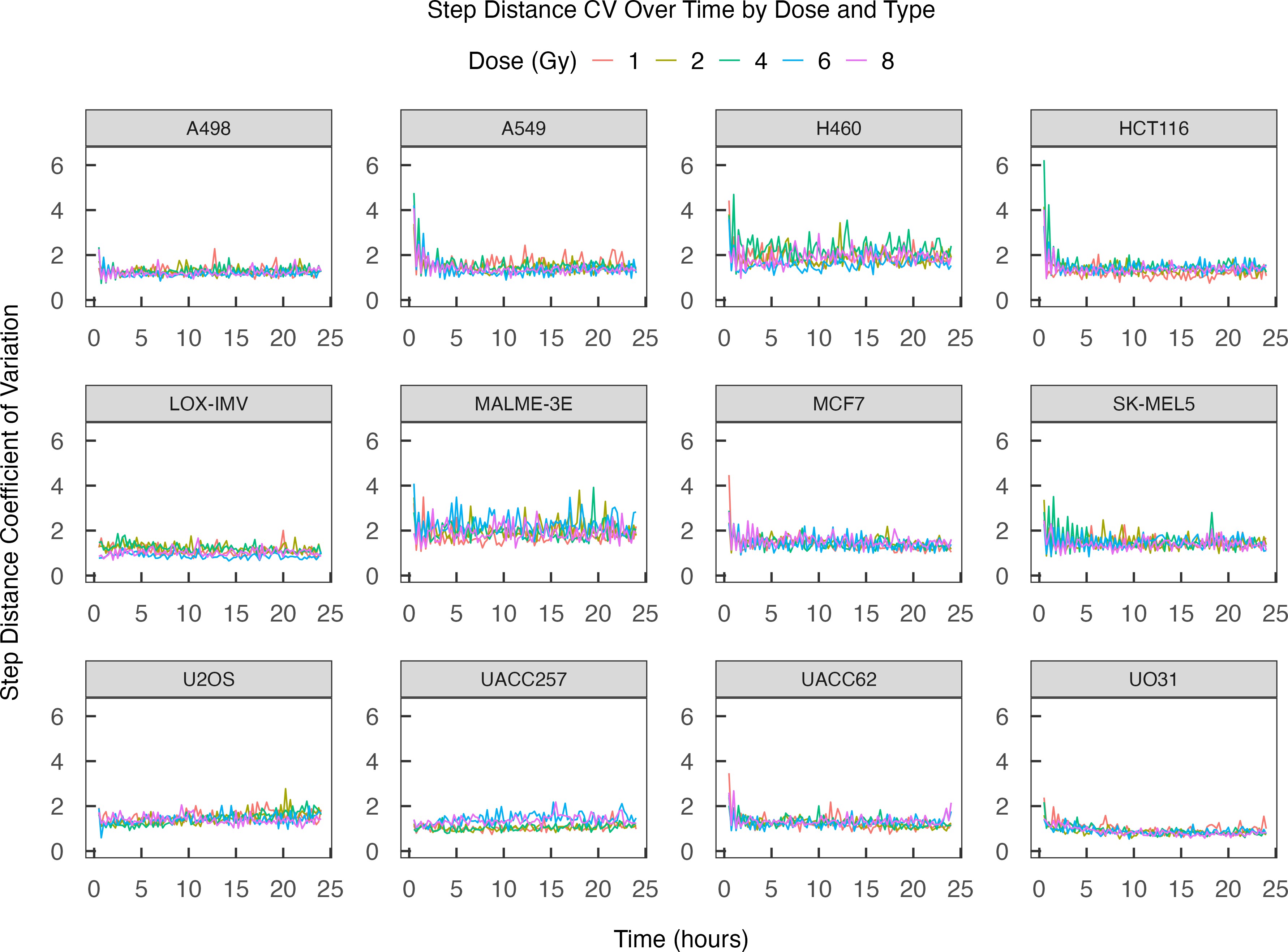

**Supplementary Fig. 2 Step distance coefficient of variation (CV) over time, grouped by radiation dose and cell line.** The migration speeds of irradiated cells remain homoscedastic (cell speeds exhibit constant variance in time), which is indicated by the largely steady CV of the 15- minute step distances over time. Cell lines A549, HCT116, MCF7, SK-MEL5, and UACC62 exhibited varying degrees of initial high variability in step length, which subsequently decreased and stabilized to a constant variance over the following hours; all cell lines exhibited relatively consistent variability over time throughout most of the experiment period.

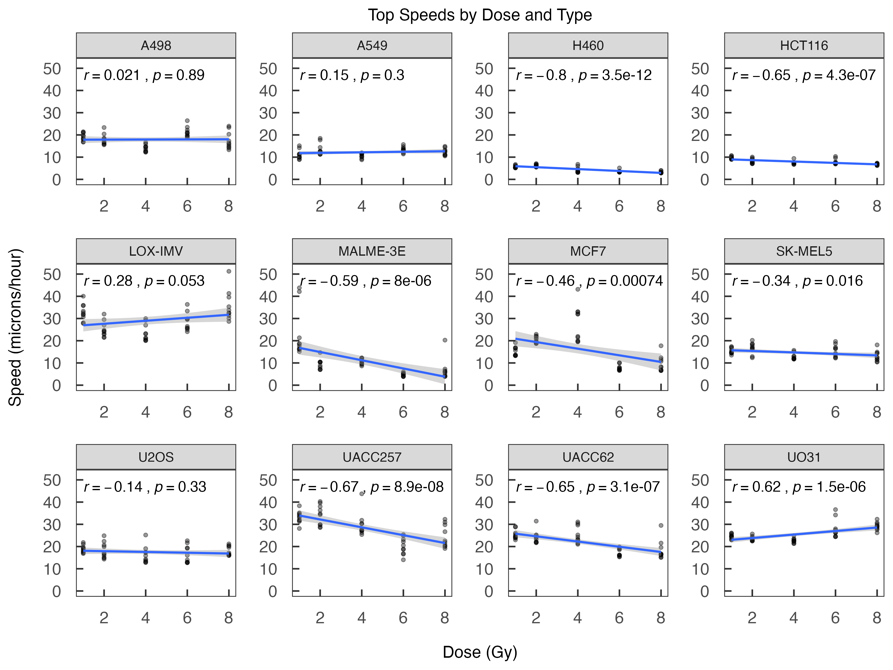

**Supplementary Fig. 3 Speeds of fastest cells irradiated at varying dose levels grouped by cell line.** We regressed the speed of the top ten fastest-moving cells against radiation dose. The Pearson correlation coefficients between dose and speed and their p-values were calculated and displayed, along with regression lines of speed on dose. No significant positive dose-speed correlation was detected in the most migratory cells from lines with non-significant or negative dose-speed correlations.

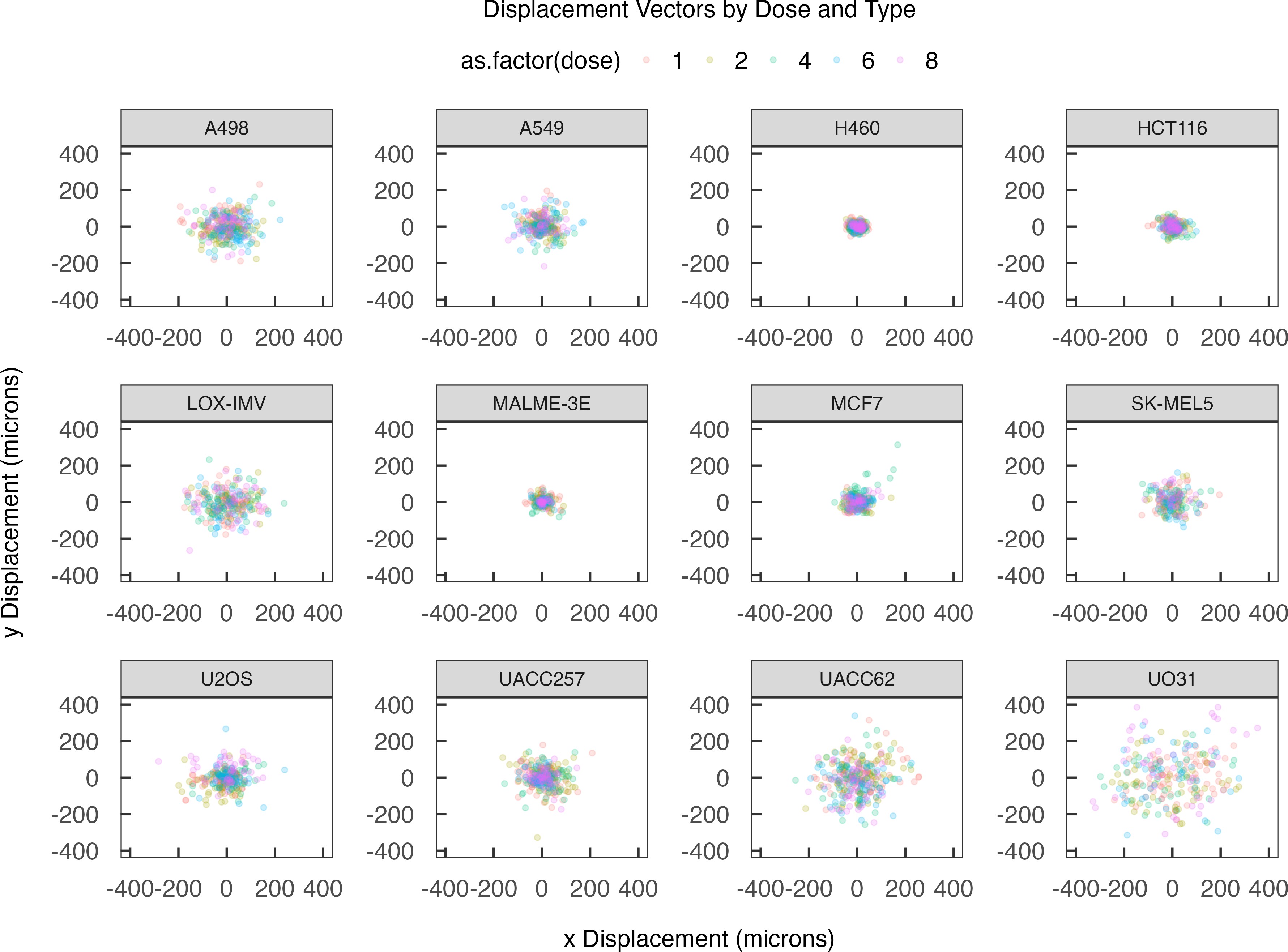

**Supplementary Fig. 4** **Displacement vectors of cells grouped by radiation dose and cell line.** The 2D displacements of cells between their initial and final positions are visualized, revealing varying degrees of dissemination in space across cell lines, with UACC62 and UO31 cells among the most dispersive. However, no directional preference was observed, indicating an absence of collective motion at the population level.

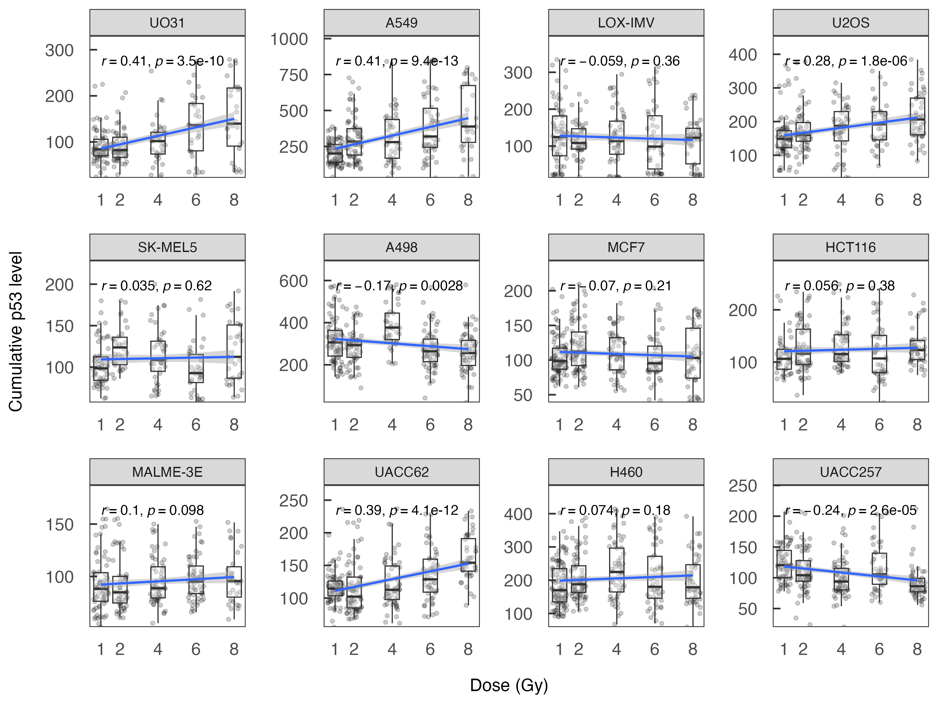

**Supplementary Fig. 5** **Cumulative p53 levels irradiated at varying dose levels grouped by cell line.** We regressed cumulative p53 level against radiation dose and calculated the Pearson correlation coefficients and their p-values. Among the 12 cell lines, UO31, A549, U2OS, and UACC62 cells exhibited significant positive correlations between cumulative p53 level and dose.

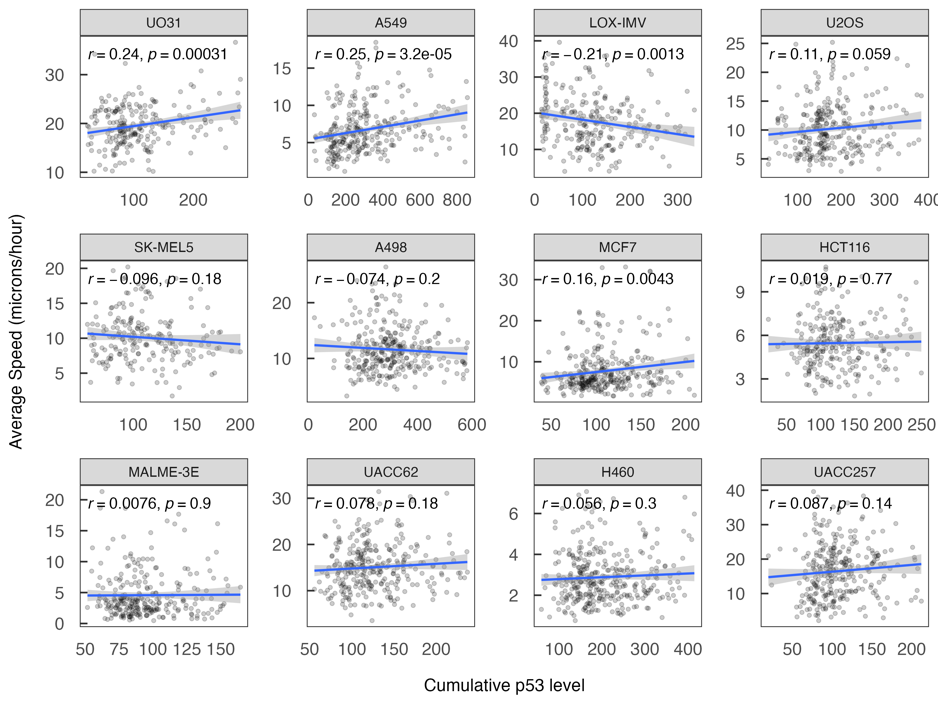

**Supplementary Fig. 6** **Correlation between average cell speed and cumulative p53 level, grouped by cell line.** We regressed average cell speed against cumulative p53 level and calculated the Pearson correlation coefficient and their p-values. UO31, A549, and MCF7 cells showed significant positive correlations between speed and cumulative p53 level.

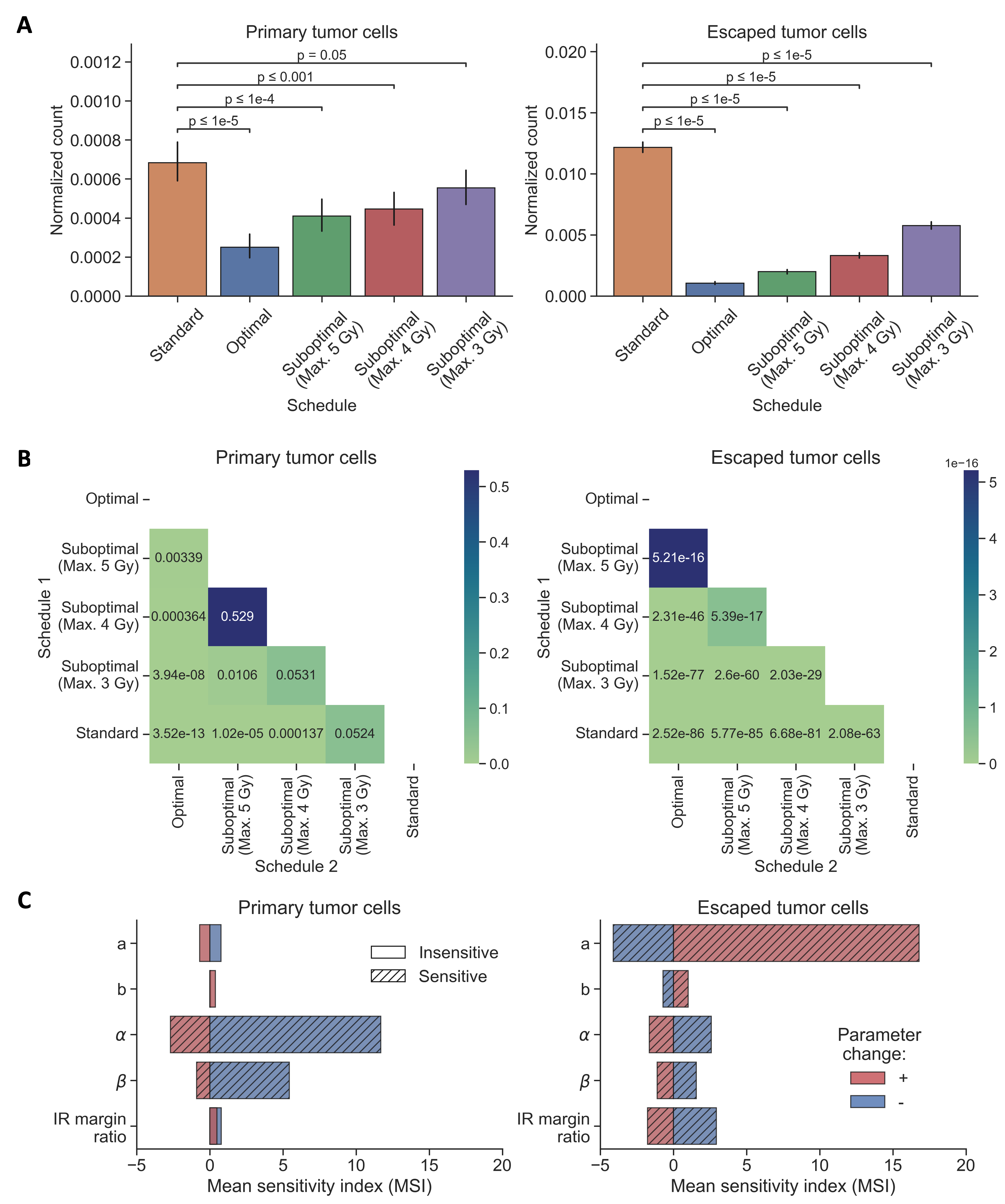

**Supplementary Fig. 7 Statistical and sensitivity analyses of model predictions of radiation response. (A)** Comparison of normalized counts of cells in primary tumor and escaped cells across different schedules from Fig. 4A. **(B)** P-values of Mann-Whitney U test to compare the schedules in panel A. **(C)** Sensitivity analyses quantifying the effect of marginal changes in the model parameters (baseline speed a, dose-speed coefficient b, linear radiosensitivity parameter α, and quadratic radiosensitivity parameter β) on the expected normalized cell count in the primary tumor and on the expected normalized number of escaped cells. Mean sensitivity indices were obtained using a derivative-based method where the parameter values were varied by up to 40% (Methods). Univariate linear regressions of treatment outcomes (PTCs and ETCs) against perturbed model parameters were performed, and statistically significant parameters at the 5% level were identified to be sensitive and displayed in a hatched pattern (see Methods).

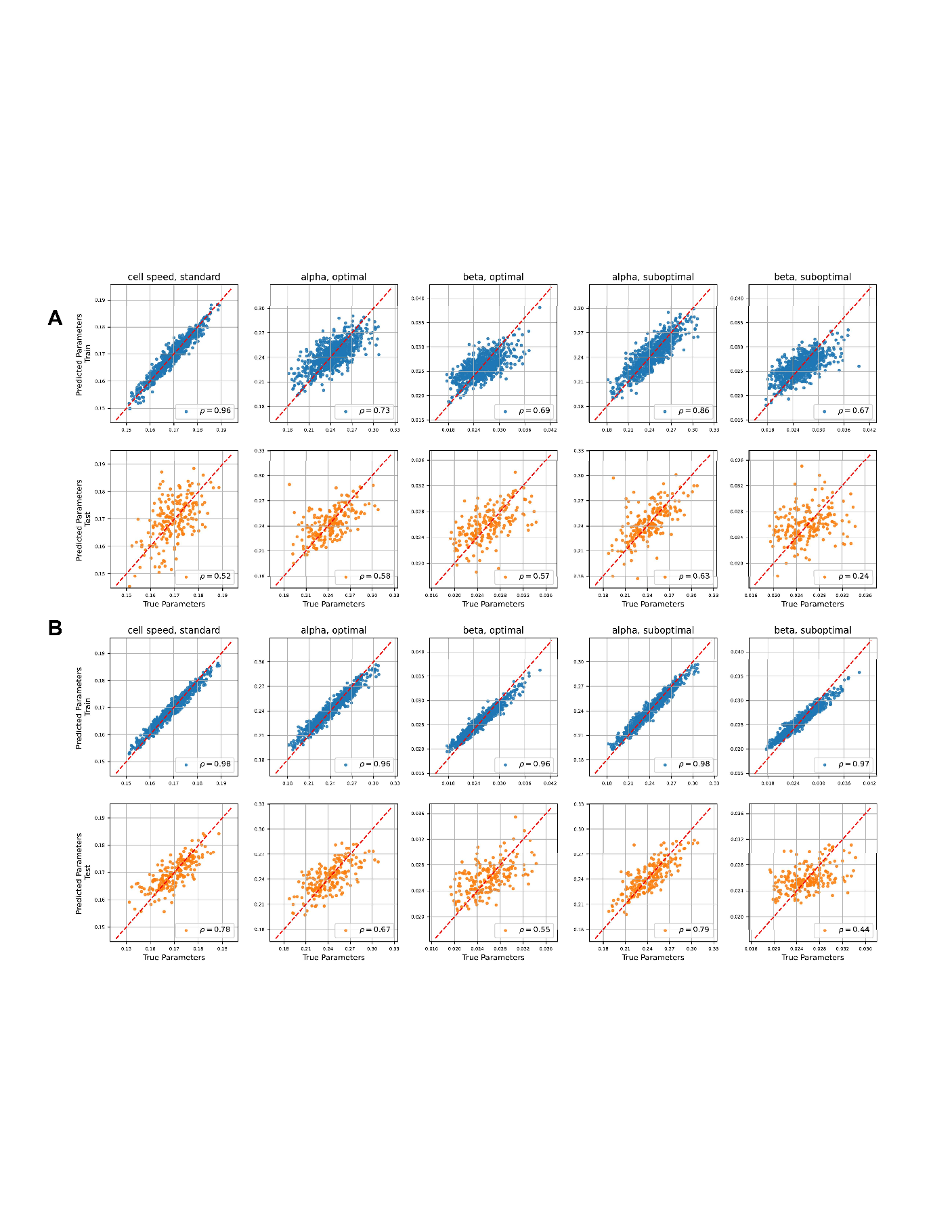
 **Supplementary Fig. 8** **Train and test distributions for baseline ML models.** **(A)** Results for Sparse Ridge Linear Regression model. **(B)** Results for random forest model. In both panels, each column represents different parameter and schedule combinations. The first row (blue dots) shows results for the 800 training trajectories, while the second row (orange dots) presents predictions for the 200 unseen test trajectories. Each scatter plot shows the relationship between true (x-axis) and predicted parameter values (y-axis). The red dashed line indicates perfect prediction, quantified with the Pearson correlation ρ, with higher values indicating stronger prediction accuracy. In both models, train distributions (blue) are narrower than test distributions (orange), indicating overfitting, although hyperparameter tuning was performed in the inner cross-validation loop. This is a common issue with traditional Machine Learning methods. On the other hand, the results for the pretrained transformer model presented in Figure 7 show the best test prediction accuracy (Fig. 7, panel B) while maintaining the most similar train and test distributions (Fig. 7, panel C), suggesting strong prediction power and out-of-distribution generalizability for unseen trajectories.

| **Cell line** | **Variable** | **Coefficient** | **P-value** |
| --- | --- | --- | --- |
| UO31 | Dose | 0.52 | 0.0026 |
|  | Cumulative p53 level $\times$ Dose | 0.0013 | 0.15 |
| A549 | Dose | 0.0042 | 0.97 |
|  | Cumulative p53 level $\times$ Dose | 0.00041 | 0.048 |

**Supplementary Tab. S1** **Linear regression model to assess the effects of radiation dose and cumulative p53 level on motility changes in UO31 and A549 cells.** We regressed average cell speed against dose and the interaction between dose and cumulative p53 level. The interaction between dose and cumulative p53 level had a significant impact on the speed of A549 cells and no significant impact on that of UO31 cells.

1. The case of zero irradiation being optimal is of course biologically unrealistic. Formally, we can rule this effect out by assuming that the tumor grows absent treatment, i.e. that *γ*(*s*) *> s >* 0 at a dose of zero.
2. We have

$$0=\frac{d}{ds}\left( \frac{s}{s-\gamma\left( s \right)} \right)=\left( \frac{s\gamma^{'}\left( s \right)-\gamma(s)}{\left( s-\gamma\left( s \right) \right)^{2}} \right)$$

   and *s* ≠ *γ*(*s*) per assumption
3. The second order condition looks as follows:

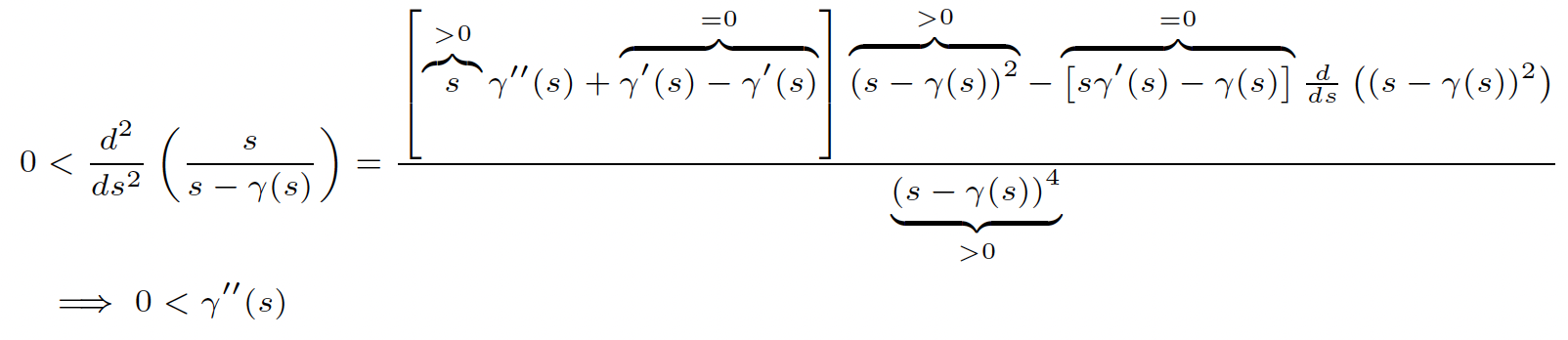
